# Thermodynamic Diversity in Staphylococcal Nitrate Sensors with a Conserved Binding Pocket

**DOI:** 10.64898/2026.09.22.751059

**Authors:** Elizabeth K. Pack, Ke Ji, Jonathan Martin, Weicheng Peng, Mariah A. Cook, Gabriele Meloni, Sheel C. Dodani

## Abstract

Nitrate is an important oxyanion in bacterial physiology, with major roles as a nitrogen source for assimilation and as an alternative electron acceptor in anaerobic respiration. Accordingly, diverse nitrate-responsive regulatory systems sense nitrate availability and control the expression of genes involved in nitrate transport and metabolism. In the NreABC system, the soluble GAF-like domain protein NreA functions as a cellular nitrate sensor. To date, our molecular understanding of nitrate recognition by NreA has been derived primarily from studies of the *Staphylococcus carnosus* protein, with less known about homologues from other species. Here, we identified 736 NreA homologues through bioinformatic analysis of the GAF-like domain superfamily and selected five staphylococcal representatives for thermodynamic characterization by isothermal titration calorimetry. Despite conservation of the nitrate-binding pocket, these homologues resolved into high- and low-affinity groups with distinct thermodynamic profiles. Comparative structural modeling and chimeragenesis of a low-affinity homologue revealed that the C-terminal region is a non-coordinating determinant of nitrate-binding affinity. Together, these findings show that a conserved nitrate-binding pocket preserves recognition, while distal sequence variation shapes thermodynamics and tunes the nitrate concentration range over which NreA sensors can function.

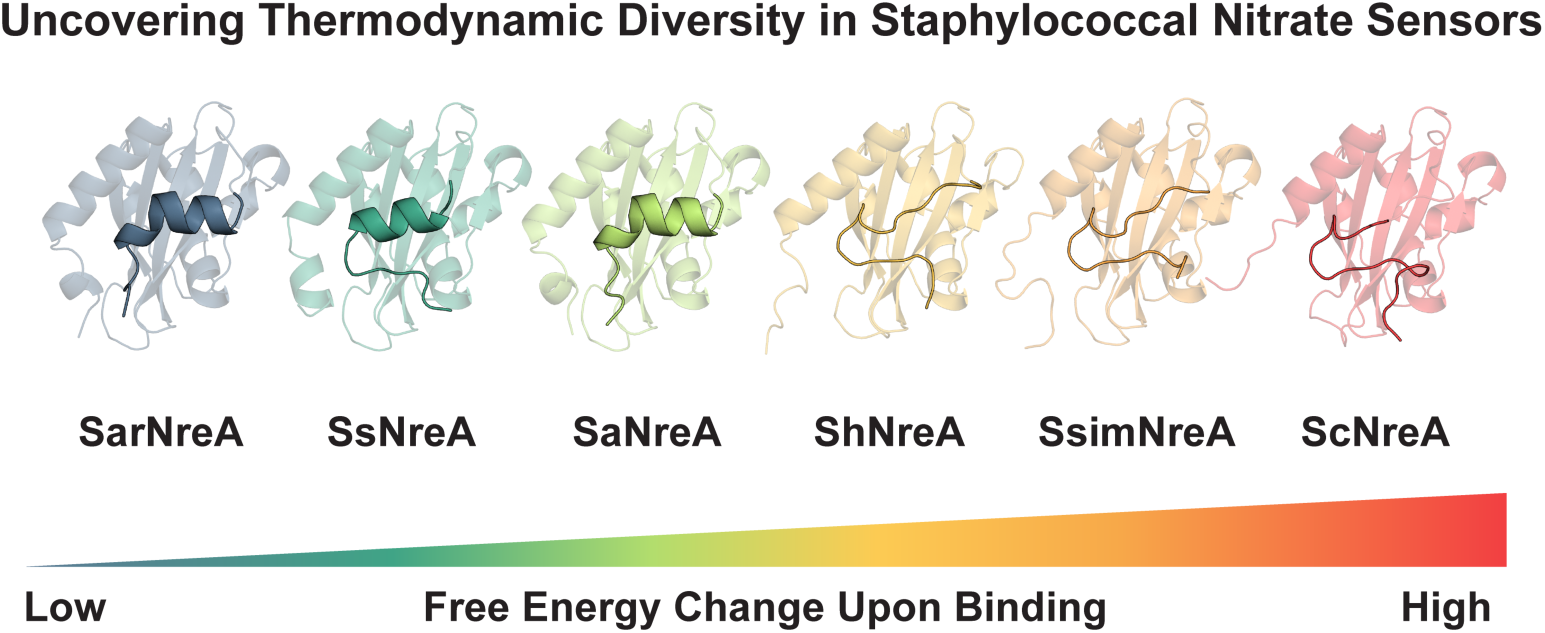

## Introduction

Nitrate is a key oxyanion in bacterial physiology, serving fundamental roles in nitrogen assimilation and anaerobic respiration (Cole & Richardson, 2008; Luque-Almagro et al., 2011; Maiti et al., 2025; Unden & Klein, 2021). Beyond these core metabolic functions, nitrate can also influence processes such as chemotaxis, biofilm formation, and host colonization (Ganusova et al., 2024; Liu et al., 2023; Mashruwala et al., 2017; Martín-Mora et al., 2019; Van Alst et al., 2007; Yang et al., 2025). As environmental conditions and metabolic demands fluctuate, bacteria must sense nitrate availability and adapt accordingly (Uehara et al., 2026; Zhang et al., 2022). At the molecular level, this response is coordinated by diverse regulatory systems that connect nitrate recognition to the expression of genes involved in nitrate transport and metabolism (Durand & Guillier, 2021).

Among these, two-component systems are widespread across bacteria and typically consist of a sensor histidine kinase that detects an environmental signal and a response regulator that mediates the downstream response (Stock et al., 2000). For example, in *Escherichia coli*, NarX and NarQ are membrane-associated nitrate-sensing histidine kinases that signal through the response regulators NarL and NarP, respectively (Durand & Guillier, 2021). In contrast, the NreABC system of *Staphylococcus carnosus* separates nitrate recognition from the histidine kinase (Schlag et al., 2008). The soluble GAF-like domain protein NreA (ScNreA) senses nitrate, whereas NreB functions as the oxygen-responsive histidine kinase and NreC as the response regulator (Fedtke et al., 2002; Niemann et al., 2014; Nilkens et al., 2014). In the absence of nitrate, ScNreA associates with NreB and inhibits NreC phosphorylation; nitrate binding weakens this interaction, enabling NreC-dependent activation of the *narGHJI* nitrate reductase operon (Fedtke et al., 2002; Klein et al., 2020). This separation has made ScNreA a tractable system for investigating the molecular basis of nitrate recognition by a soluble sensor protein.

The X-ray crystal structure of nitrate-bound ScNreA revealed key features of nitrate recognition (Niemann et al., 2014). ScNreA has a hydrophobic binding pocket within an electropositive environment created by the α3 and α6 helix dipoles (Figure 1A; Figure S1) (Niemann et al., 2014). Specifically, nitrate is coordinated by the indole N-H of W45 on β2 and the backbone N-H of L67 and A68 on *a*3 and I97 on *a*6 (Niemann et al., 2014). Additional residues, including L61, G66, Y95, and P96, line the pocket, with Y95 proposed to function as a lid (Niemann et al., 2014). Structure-guided mutagenesis linked nitrate-binding affinity to the sensory function of ScNreA. Relative to the wild type (*K*_d_ ≈ 22 µM in 200 mM NaCl), the W45F and Y95L variants had weaker affinities (*K*_d_ ≈ 62 µM and *K*_d_ ≈ 61 µM, respectively) but retained nitrate-responsive regulation (Niemann et al., 2014). In contrast, the Y95A variant bound nitrate more weakly (*K*_d_ ≈ 106 µM) and lost nitrate-responsive regulation (Niemann et al., 2014). These findings led to the proposal of an affinity threshold for functional nitrate sensing by ScNreA (Niemann et al., 2014).

**Figure 1.**
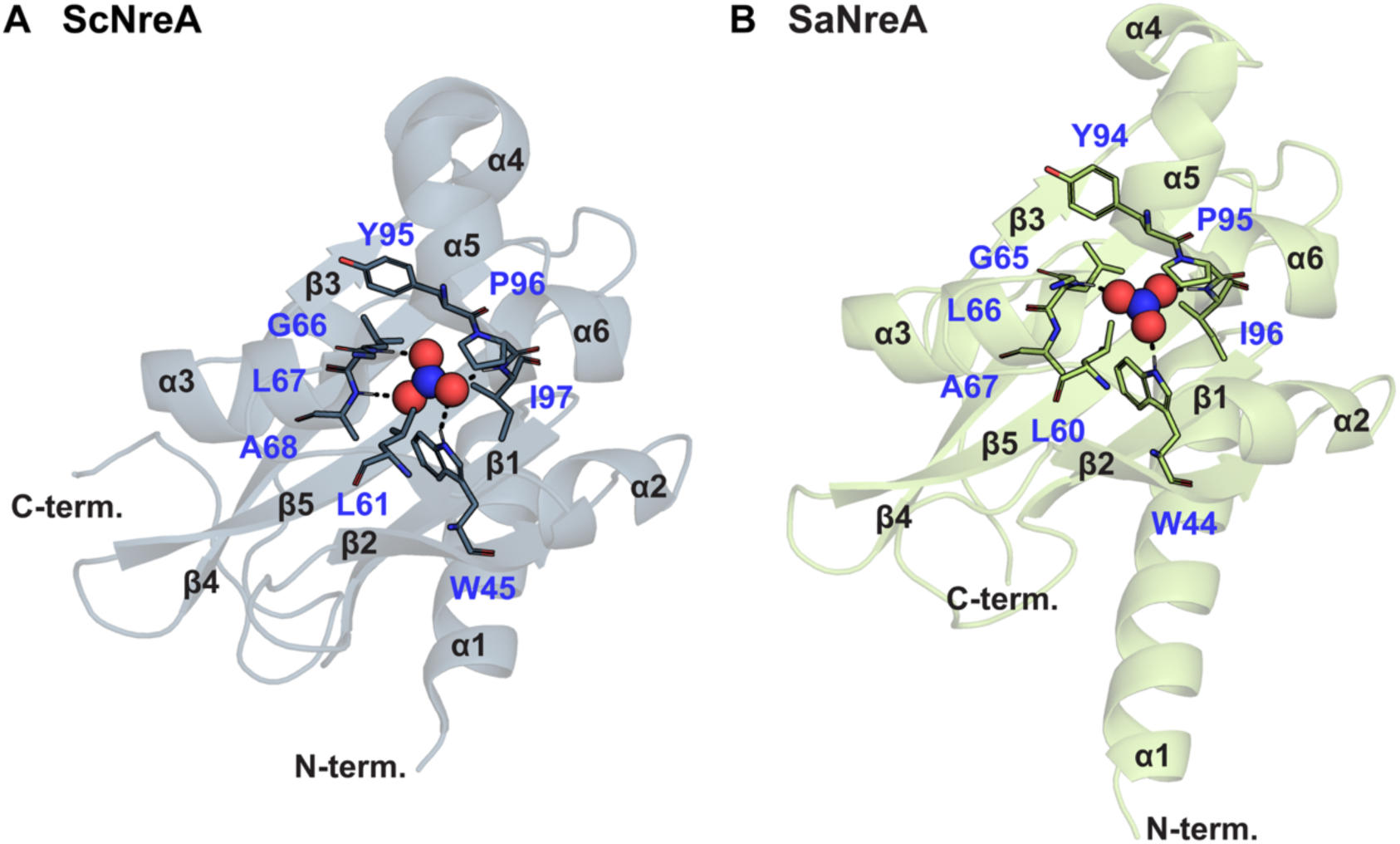
X-ray crystal structures of nitrate-bound NreA from (A) *Staphylococcus carnosus* (ScNreA, PDB ID: 4IUK) and (B) *Staphylococcus aureus* (SaNreA, PDB ID: 6IZJ). All residues within 4 Å of the bound nitrate in the ScNreA structure are shown and labeled for both panels. All hydrogen-bonding interactions are indicated by dashed lines. The α-helices and β-strands are labeled according to the structural nomenclature established for ScNreA (Niemann et al., 2014). The N- and C-termini are labeled. Residues 1–10 on the N-terminus and 154–155 on the C-terminus of ScNreA; residues 1–3 on the N-terminus and 142–150 on the C-terminus of SaNreA were not resolved in the structures.

Subsequent structural characterization of a homologous NreA from *S. aureus* (SaNreA) showed conservation of the overall fold and nitrate-binding pocket (Figure 1B; Figure S1) (Sangare et al., 2020). Comparison of the nitrate-bound SaNreA with the corresponding Y94A apo-like mutant structure further revealed conformational variability in the C-terminal region, distal to the nitrate-binding pocket (Figure S1) (Sangare et al., 2020). The C-terminus adopts a loop-like conformation in the nitrate-bound SaNreA but forms an α-helix in the Y94A apo-like mutant (Sangare et al., 2020). This rearrangement alters the local surface electrostatic potential and has been proposed to modulate the NreA-NreB interaction (Figure S2) (Sangare et al., 2020). This distal conformational variability raised the possibility that regions outside the nitrate-binding pocket could contribute to nitrate recognition and downstream NreB/NreC signaling.

More recently, our thermodynamic, kinetic, and computational analyses of ScNreA have further defined the basis of nitrate recognition and anion selectivity (Ji et al., 2025). Together, these studies have provided a molecular understanding of nitrate sensing by ScNreA, but whether nitrate-binding thermodynamics vary across NreA homologues and what structural features contribute to such variation remain unclear. Here, we used bioinformatic analysis to identify ScNreA homologues and selected a set of staphylococcal representatives with conserved nitrate-binding pocket residues for thermodynamic characterization by isothermal titration calorimetry (ITC). We then combined comparative structural modeling with C-terminal chimeragenesis to test whether a distal, non-coordinating region contributes to nitrate recognition.

## Methods

### General

All reagents and supplies used in this study were acquired from Sigma Aldrich (St. Louis, Missouri), Thermo Fisher Scientific (Waltham, Massachusetts), Thomas Scientific (Swedesboro, New Jersey), VWR (Radnor, Pennsylvania), USA Scientific (Ocala, Florida), or Research Products International (Mount Prospect, Illinois), except where noted. PyMOL v3.1.6.1 was used to render and visualize all X-ray crystal structures and AlphaFold3 models (Abramson et al., 2024; The PyMOL Molecular Graphics System).

### Bioinformatic Pipeline to Identify ScNreA Homologues

The sequences in the GAF-like domain superfamily (IPR029216) were downloaded from the InterPro database on July 22, 2023 (Zenodo File 1) and filtered using Python to retain sequences between 100 and 200 amino acids in length (Zenodo File 2, 3) (Blum et al., 2025). The filtered sequence set was searched using phmmer in HMMER 3.3.2, with the NreA sequence from *S*. *carnosus* (ScNreA, UniProt ID: B9DL91) as the query (Eddy, 2011). An E-value of 1 × 10^-3^ was used to identify sequences homologous to ScNreA (Zenodo File 4). The resulting phmmer-generated query-to-hit sequence alignments were then used to identify positions corresponding to the ScNreA nitrate-binding pocket residues W45, L67, A68, Y95, and I97, and the conservation of these residues was analyzed using Python (Zenodo File 5– 7). Finally, all non-staphylococcal sequences and redundant sequences from the same staphylococcal species were removed to generate the final set of putative NreA sequences (Table S1).

Next, the Enzyme Function Initiative-Enzyme Similarity Tool (EFI-EST) and EFI-Genome Neighborhood Tool (EFI-GNT) were used to evaluate the genes upstream and downstream of the NreAs in the final sequence set (Oberg et al., 2023). A Sequence Similarity Network (SSN) was generated using the EFI-EST for the GAF-like domain superfamily with the following parameters: i) UniRef50 clusters were selected, ii) UniProt-defined fragment sequences were excluded, iii) a default fraction of 1 was used, iv) an E-value of 5 was used, and v) an alignment threshold score of 35 was used to finalize the SSN. The resulting SSN was analyzed using the EFI-GNT to generate a Genome Neighborhood Diagram (GND) with a neighborhood size of 20 and a minimum co-occurrence threshold of 20% (Figure S3).

After genome neighborhood analysis, five sequences were selected for this study based on theoretical isoelectric point (pI) and percent sequence identity relative to ScNreA and SaNreA. The theoretical pI was determined using the ExPASy ProtParam tool, and the percent sequence identity was determined using the EMBL-EBI Clustal Omega MSA tool (Table S1; Figure S4) (Bateman et al., 2025; Gasteiger et al., 2005; Sievers et al., 2011). The putative NreAs included *S*. *simulans* (SsimNreA; UniProt ID: A0A855LGD0), *S*. *haemolyticus* (ShNreA; strain JCSC1435; UniProt ID: Q4L8Q8), *S. argenteus* (SarNreA; UniProt ID: A0A7U7PWK7), *S. schleiferi* (SsNreA; UniProt ID: A0A7Z7W3U1), and *S*. *aureus* (SaNreA; strain NCTC 8325/PS47, UniProt ID: Q2FVM5) (Bateman et al., 2025). The AlphaFold3 web server (https://alphafoldserver.com) was used to generate all models (Figure S5, S6) (Abramson et al., 2024).

### Plasmid Design and Preparation

The ScNreA plasmid was generated in our previous study (Ji et al., 2025). Like ScNreA, all genes were codon optimized for expression in *Escherichia coli* K12 and cloned into the pET-21a(+) vector between the NdeI and HindIII restriction sites in frame with the vector-encoded C-terminal polyhistidine tag (GenScript, Piscataway, New Jersey, Figure S7–S13).

The commercially prepared plasmids were resuspended in 20 µL autoclaved water to a final concentration of 5 ng/µL. *E. cloni* 10G ELITE Competent Cells (Lucigen, Middleton, Wisconsin) were transformed with 5 ng of each plasmid using electroporation (Bio-Rad Laboratories, Hercules, California). The resulting transformation mixture was diluted with 750 µL of pre-warmed Super Optimal broth with Catabolite repression (SOC) medium and incubated at 37 °C for 45 min with shaking at 250 rpm (LSE Benchtop Shaking Incubator, Corning, Corning, New York). After incubation, the transformation mixture was plated onto a Miller’s Luria Broth (LB) agar with 100 µg/mL ampicillin and incubated for ∼16 h at 37 °C (12-140 Incubator, Quincy Lab, Inc., Burr Ridge, Illinois). A single colony was inoculated into 5 mL of LB medium with 100 µg/mL ampicillin and incubated for ∼16 h at 37 °C with shaking at 250 rpm (New Brunswick Innova 42R, Eppendorf, Enfield, Connecticut), followed by harvesting by centrifugation at 18,000*g* for 15 min at 20 °C (5424R, Eppendorf, Enfield, Connecticut). The plasmid was isolated from the resulting cell pellet using the QIAprep Spin Miniprep Kit (Qiagen) according to the manufacturer’s instructions.

### Molecular Cloning and Preparation of SaNreA Chimera

The gene encoding the SaNreA chimera was generated using the ScNreA (insert) and SaNreA (backbone) plasmids (Figure S14). One primer set was designed to amplify a fragment encoding the C-terminal 16 amino acids of ScNreA (I140–P155), with a complementary overlap to the SaNreA backbone fragment (Table S2). The second primer set was designed to amplify the pET-21a(+) vector with SaNreA without its C-terminal sixteen amino acids (D135– K150), with a complementary overlap to the amplified ScNreA fragment (Table S2). Each primer was resuspended and diluted in autoclaved water to a final concentration of 10 µM. Both templates were diluted in autoclaved water to a final concentration of 10 ng/µL. Polymerase chain reaction (PCR) amplification was carried out using Phusion Hot Start Flex 2X Master Mix (New England Biolabs, Ipswich, Massachusetts) according to the conditions listed in Table S3 (Bio-Rad T100 Thermal Cycler, Bio-Rad, Hercules, California). After the PCR, each reaction mixture was treated with 1 µL of DpnI (New England Biolabs, Ipswich, Massachusetts) and incubated for 2 h at 37 °C (Bio-Rad T100 Thermal Cycler, Bio-Rad, Hercules, California).

Next, the PCR products were separated by electrophoresis on a 1% agarose gel (Gold Biotechnology, St. Louis, Missouri) in 1X Tris-Acetate-EDTA (TAE) running buffer for 20 min at 110 V using a Mini-Sub Cell GT and PowerPac (Bio-Rad Laboratories, Hercules, California). The DNA was then extracted using the Zymoclean Gel DNA Recovery Kit (Zymo Research, Irvine, California) and quantified using a NanoDrop Lite spectrophotometer. For the assembly, 2.5 ng of insert and 100 ng of backbone were combined with 10 µL of NEBuilder HiFi DNA Assembly Master Mix (New England Biolabs, Ipswich, Massachusetts) and incubated for 1 h at 50 °C. The assembled DNA was purified using the Zymo DNA Clean and Concentrator 5 kit (Zymo Research, Irvine, California). For sequence verification by Sanger sequencing (Eurofins, Luxembourg City, Luxembourg), a portion of the DNA was diluted to 5 ng/µL, transformed into *E. cloni* 10G ELITE Competent Cells, and isolated as described in the *Plasmid Design and Preparation* section.

### Protein Expression and Purification

The entire procedure in this section was carried out twice to generate two independent protein preparations for SsimNreA, ShNreA, SarNreA, and SsNreA. For SaNreA and the SaNreA chimera, the procedure was carried out four times to generate four independent protein preparations, with two used for ITC and two used for circular dichroism. All plasmids were freshly transformed into *E. cloni* 10G ELITE Competent Cells, isolated as described in the *Plasmid Design and Preparation section* and subsequently transformed into *E. cloni* EXPRESS BL21 (DE3) Electrocompetent Cells (Lucigen, Middleton, Wisconsin).

The following procedure was adapted from our previous study to express SsimNreA, ShNreA, and the SaNreA chimera (Ji et al., 2025). For each protein, a single colony of the transformed *E. cloni* EXPRESS BL21 (DE3) Competent Cells was inoculated into 30 mL of Terrific Broth (TB) medium with 100 µg/mL ampicillin and incubated for ∼13 h at 30 °C with shaking at 230 rpm (New Brunswick Innova 42R, Eppendorf, Enfield, Connecticut). Four 750 µL aliquots of the overnight culture were used to inoculate four 600 mL TB flasks with 100 µg/mL ampicillin and incubated at 37 °C with shaking at 250 rpm until the optical density at 600 nm (OD_600_) reached ∼5. The cultures were cooled to 18 °C over 1 h with shaking at 250 rpm, and then 300 µL of 1 M isopropyl β-D-1-thiogalactopyranoside (IPTG) was added to each flask to give a final concentration of 0.5 mM and induce protein expression. The cultures were incubated for an additional 20 h at 18 °C with shaking at 250 rpm. The next day, the cells were harvested by centrifugation at 3,000*g* for 45 min at 4 °C (5810 R, Eppendorf, Enfield, Connecticut), and resuspended in pre-chilled 50 mM HEPES buffer (pH 7.8) with 200 mM NaCl and reharvested by centrifugation at 3,000*g* for 25 min at 4 °C. The final cell pellet was stored at -20 °C for no longer than one week before purification.

The following procedure was adapted from a previously reported procedure for SaNreA for the expression of SarNreA, SsNreA, and SaNreA (Sangare et al., 2020). For SarNreA and SsNreA, a single colony of the transformed *E. cloni* EXPRESS BL21 (DE3) Competent Cells was inoculated into 30 mL of LB medium with 100 µg/mL ampicillin and incubated for ∼16 h at 30 °C with shaking at 230 rpm (New Brunswick Innova 42R, Eppendorf, Enfield, Connecticut). For SaNreA, a single colony of the transformed *E. cloni* EXPRESS BL21 (DE3) Competent Cells was inoculated into 30 mL of LB medium with 100 µg/mL ampicillin and incubated for ∼13 h at 30 °C with shaking at 230 rpm (New Brunswick Innova 42R, Eppendorf, Enfield, Connecticut). Four 6 mL aliquots of the overnight culture were used to inoculate four 600 mL LB flasks with 100 µg/mL ampicillin and incubated at 37 °C with shaking at 250 rpm until the OD_600_ reached between ∼0.6–0.8 for SarNreA and SsNreA and ∼0.8–1 for SaNreA. Then, 240 µL of 1 M IPTG was added to each flask to give a final concentration of 0.4 mM and to induce protein expression. The cultures were incubated for an additional 4 h at 30 °C with shaking at 250 rpm followed by harvesting and storage as described above.

For each protein preparation, the frozen cell pellet was first thawed in an ice water bath and then resuspended in 4 mL of pre-chilled lysis buffer (20 mM Tris buffer [pH 7.5 at 4 °C] with 200 mM NaCl, 5 mM magnesium chloride, and 30 µg/mL deoxyribonuclease I) per gram of wet cell pellet by vortexing. The resuspended cell pellet was lysed by sonication in an ice-water bath with a 30% amplitude setting for 5 min using cycles of 15 sec on and 45 sec off (QSonica, Newton, Connecticut). The cell lysate was clarified by centrifugation at 30,000*g* for 35 min at 4 °C (Optima XPN-80 Ultracentrifuge, Beckman Coulter, Brea, California).

For all chromatography described below, an NGC Quest 10 Chromatography System with a sample pump loading system (Bio-Rad Laboratories, Hercules, California) was used in a refrigerator maintained between 4 °C and 15 °C. For affinity purification, prior to loading the sample, a 5 mL HisTrap HP column pre-packed with a Nickel Sepharose High Performance (HP) resin (Cytiva, Marlborough, Massachusetts) was pre-equilibrated with 5 column volumes (CV) of buffer A (20 mM Tris buffer at pH 7.5, 15 °C with 200 mM NaCl and 30 mM imidazole). The clarified supernatant was loaded onto the column at a flow rate of 4 mL/min, and then the column was washed at a flow rate of 4 mL/min with 20 CV of buffer A. Following this step, the proteins were eluted at a flow rate of 4 mL/min of buffer A using a 0–100% gradient of buffer B (20 mM Tris buffer [pH 7.5 at 15 °C] with 200 mM NaCl and 500 mM imidazole) over 17 CV for all proteins except the SaNreA chimera, which required 23 CV. All fractions eluted during the gradient that had an absorbance at 280 nm were combined for desalting.

For desalting, a 50 mL HiPrep 26/10 desalting column (Cytiva, Marlborough, Massachusetts) was pre-equilibrated with 5 CV of buffer C (50 mM HEPES buffer [pH 7.8 at 10 °C] with 200 mM NaCl). The pooled protein fractions were loaded at a flow rate of 5 mL/min, and the protein was eluted at a flow rate of 4 mL/min with buffer C over 2 CV. All fractions with an absorbance at 280 nm were combined and concentrated using a 15 mL Millipore Amicon Ultra Centrifugal Filter Tube with a 10 kDa molecular weight cut-off (MWCO) by centrifugation at 2,500*g* at 4 °C (5424 R, Eppendorf, Enfield, Connecticut) for size-exclusion chromatography (SEC). To prevent protein precipitation, the pooled fractions of SsimNreA, ShNreA, and the SaNreA chimera were concentrated to a maximum of 300 µM and SaNreA, SsNreA, and SarNreA were concentrated to a maximum of 250 µM. The concentrations were estimated during centrifugation using a NanoDrop Lite Spectrophotometer.

For SEC, a HiLoad 26/600 Superdex 200 size-exclusion column (Cytiva, Marlborough, Massachusetts) was pre-equilibrated with 2 CV of buffer C. The pooled fractions were loaded at a flow rate of 2.6 mL/min, and the protein was eluted at a flow rate of 2.6 mL/min with buffer C over 1 CV, the same elution buffer used for desalting (Figure S15–S20). All fractions with an absorbance at 280 nm corresponding to a monomeric protein based on comparison with Gel Filtration Standard #1511901 (Bio-Rad Laboratories, Hercules, California) were combined and concentrated to ∼10 mL using a 15 mL Millipore Amicon Ultra Centrifugal Filter Tube with a 10 kDa MWCO at 2,500*g* at 4 °C (5424 R, Eppendorf, Enfield, Connecticut). Once concentrated, ∼10 mL of protein sample was loaded into a Slide-A-Lyzer Dialysis Cassette with a 10 kDa MWCO and dialyzed against 1 L of ITC Buffer (50 mM HEPES buffer [pH 7.8 at 10 °C]) with 50 mM NaCl (ITC buffer) for a total of 22 h with one buffer change after 10 h, at 4 °C–15 °C. Each dialysis step had a sample to buffer ratio (v/v) of 1:100 giving an overall dilution ratio of 1:10,000. The final protein sample was clarified with centrifugation at 18,000*g* for 15 min at 4 °C (5424 R, Eppendorf, Enfield, Connecticut) and stored at 4 °C for further characterization.

### Sodium Dodecyl Sulfate-Polyacrylamide Gel Electrophoresis (SDS-PAGE)

Following dialysis, the purity of each protein preparation was evaluated by SDS-PAGE. The concentration of each protein preparation was estimated using a NanoDrop Lite Spectrophotometer, and a portion was diluted to ∼0.1 mg/mL for further analysis. Each protein sample was combined with 4X Laemmli Sample Buffer (Bio-Rad Laboratories, Hercules, California) with 10% β-mercaptoethanol in a 3:1 (v/v) ratio for a total volume of 20 µL and heated at 95 °C for 5 min with shaking at 300 rpm (ThermoMixer C, Eppendorf, Enfield, Connecticut). For a given protein, 10 µL of each protein preparation and 4 µL of a pre-stained protein standard (PageRuler^TM^ Plus Prestained Protein Ladder, Thermo Scientific, Waltham, Massachusetts) were loaded onto a 12% TGX FastCast acrylamide gel (Bio-Rad Laboratories, Hercules, California). Electrophoresis was carried out in 1X Tris-Glycine-SDS running buffer for 1.5 h at 150 V using a Mini-PROTEAN Tetra Cell and PowerPac (Bio-Rad Laboratories, Hercules, California). After electrophoresis, the gel was rinsed in deionized water and heated in a microwave for 15 sec. This process was repeated three times. After the final rinse, the gel was stained with Coomassie Brilliant Blue R-250 (182 µM) in 7% acetic acid, 40% methanol, and 53% deionized water. The gel was heated in a microwave for 30 sec and then incubated at room temperature for 30 min with orbital shaking (Belly Dancer, IBI Scientific, Dubuque, Iowa). Following this, the gel was destained in a solution of 5% acetic acid, 20% ethanol, and 75% deionized water at room temperature for 4 h with orbital shaking. The destaining solution was replaced every hour. The final gel was visualized using an Azure 400 Gel Imaging System (Azure Biosystems, Dublin, California), and the image was processed with Image Lab software for PC Version 6.0 (Bio-Rad Laboratories, Hercules, California, Figure S15–S20).

### Isothermal Titration Calorimetry (ITC)

The following procedure was adapted from our previous study with ScNreA (Ji et al., 2025). The concentration of each protein preparation was first determined using the theoretical molar extinction coefficient calculated with the ExPASy ProtParam tool (Table S4) (Gasteiger et al., 2005). After dialysis, a 100 µL portion of each protein was diluted ten-fold in the ITC buffer (v/v), and then this solution was serially diluted in the ITC buffer five more times to give relative dilution factors of 1 (initial dilution), 0.75, 0.5, 0.25, 0.125, and 0.0625.

Three 100 µL aliquots of each dilution were transferred to a clear 96-well Half Area UV-Star microtiter plate (Greiner Bio-One, Monroe, North Carolina), and absorbance spectra were collected from 250 nm–600 nm with a 2 nm step size and 3.5 nm bandwidth. The absorbance intensity was baseline corrected by subtracting the average absorbance at 600 nm from the average absorbance at 280 nm. The corrected absorbance at 280 nm was plotted versus the dilution factor. The data were fit to a linear regression, and the slope was input into the following equation:

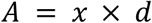

where *A* is equal to the absorbance at 280 nm, *x* is the value of the slope from the linear regression, and *d* is the initial dilution factor of 10. The absorbance (*A*) is then used to calculate the protein concentration (*c*) using the Beer-Lambert law as follows:

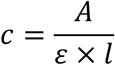

where ε is the theoretical extinction coefficient at 280 nm, and *l* is the optical pathlength (0.6 cm) (Greiner Bio-One, 2021).

ITC experiments were carried out using an Affinity ITC instrument equipped with a low-volume cell (TA Instruments, New Castle, Delaware) as described in our previous study with ScNreA (Ji et al., 2025). To minimize buffer mismatch, all sodium nitrate (NaNO_3_) solutions were prepared using the ITC buffer from the final round of dialysis. The final protein concentrations varied between protein preparations, but all NaNO_3_ solutions had a concentration 10-fold greater than the corresponding protein concentration. For each titration experiment, one 0.5 µL injection was followed by twenty-four 1.5 µL injections with a stirring rate of 125 rpm. To account for the heat of dilution, NaNO_3_ was titrated into the ITC buffer at the desired temperature. For all proteins, two independent protein preparations were evaluated in triplicate at 20 °C (Figure S21–S28; Table S5–S12). To determine the heat-capacity change (ΔC_p_), SaNreA and the SaNreA chimera were also evaluated at 10 °C, 15 °C, and 25 °C (Figure S25–S28; Table S9–S12) in triplicate with the same protein preparations used for evaluation at 20 °C.

For analysis, NITPIC was used to subtract the heat of dilution injection-by-injection from the raw data for each technical replicate before integration and baseline correction were conducted (Keller et al., 2012). Next, SEDPHAT was used to analyze the three technical measurements for each protein preparation using the global analysis tool with an “A + B ↔ AB Hetero-Association” model (Brautigam et al., 2016; Zhao et al., 2015). The best-fit parameter values for the dissociation constant (*K*_d_), enthalpy (ΔH), entropic contribution (TΔS), Gibbs free-energy change (ΔG), and the concentration correction factor were then determined for the three technical replicates for each protein preparation with the built-in error analysis using a confidence level of 0.683. To visualize the data, GUSSI was used to plot the NITPIC-generated SVD-reconstructed thermograms and the SEDPHAT-fitted isotherms with residuals (Brautigam, 2015). To obtain the heat-capacity change (ΔC_p_), three technical replicates for each temperature were evaluated for each protein preparation using an “A + B ↔ AB Hetero-Association Global Temperature Variation Analysis” model in SEDPHAT with a confidence level of 0.683. Microsoft Excel was used to average the data and calculate standard deviation across the two protein preparations.

### Circular Dichroism (CD)

CD experiments were carried out using a J-1000 Circular Dichroism Spectropolarimeter (Jasco, Easton, Maryland) in the Department of Chemistry and Biochemistry at UT Dallas. Two new protein preparations were prepared for CD experiments. The following procedure was adapted from previous studies (Greenfield, 2006; Miles et al., 2021). To reduce background absorption from the HEPES buffer, both preparations of SaNreA and SaNreA chimera were exchanged into CD buffer (10 mM phosphate buffer [pH 7.8 at 20 °C] with 50 mM NaCl) using a 15 mL Millipore Amicon Ultra Centrifugal Filter Tube with a 10 kDa MWCO by centrifugation at 2,500*g* for 50 min at 4 °C (5424 R, Eppendorf, Enfield, Connecticut) for a total of six 1:10 dilutions. ITC was carried out to correct the protein concentration as described in the *Isothermal Titration Calorimetry* section above. From this determination, each protein was diluted to 1 mg/mL in the CD buffer, and then to a final concentration of 0.1 mg/mL in the CD buffer in the absence or presence of 1 mM NaNO_3_. Prior to the start of data acquisition, 500 µL of each protein sample was incubated in a 1 mm quartz cuvette for 5 min at 20 °C in the cuvette holder. The following instrument settings were used: spectral range of 180–260 nm, 1 nm bandwidth, 20 nm/min scan speed, 0.5 nm data pitch, 1 sec response time, and 15 accumulated spectra (Figure S29, S30).

## Results

### Bioinformatic Mining for ScNreA Homologues

To identify homologues of ScNreA, we performed a bioinformatic search across the GAF-like domain superfamily (InterPro ID: IPR029216) (Figure 2), which consists of 673,187 sequences of varying lengths and domain compositions. Sequences were therefore filtered to retain those between 100 and 200 amino acids in length, consistent with ScNreA, yielding 41,540 sequences. A phmmer search using ScNreA as the query identified 736 homologous protein sequences (Eddy, 2011). Screening these sequences for conservation of W45, L67, A68, Y95, and I97 yielded 75 sequences in which all five residues were conserved (Zenodo File 7).

**Figure 2.**
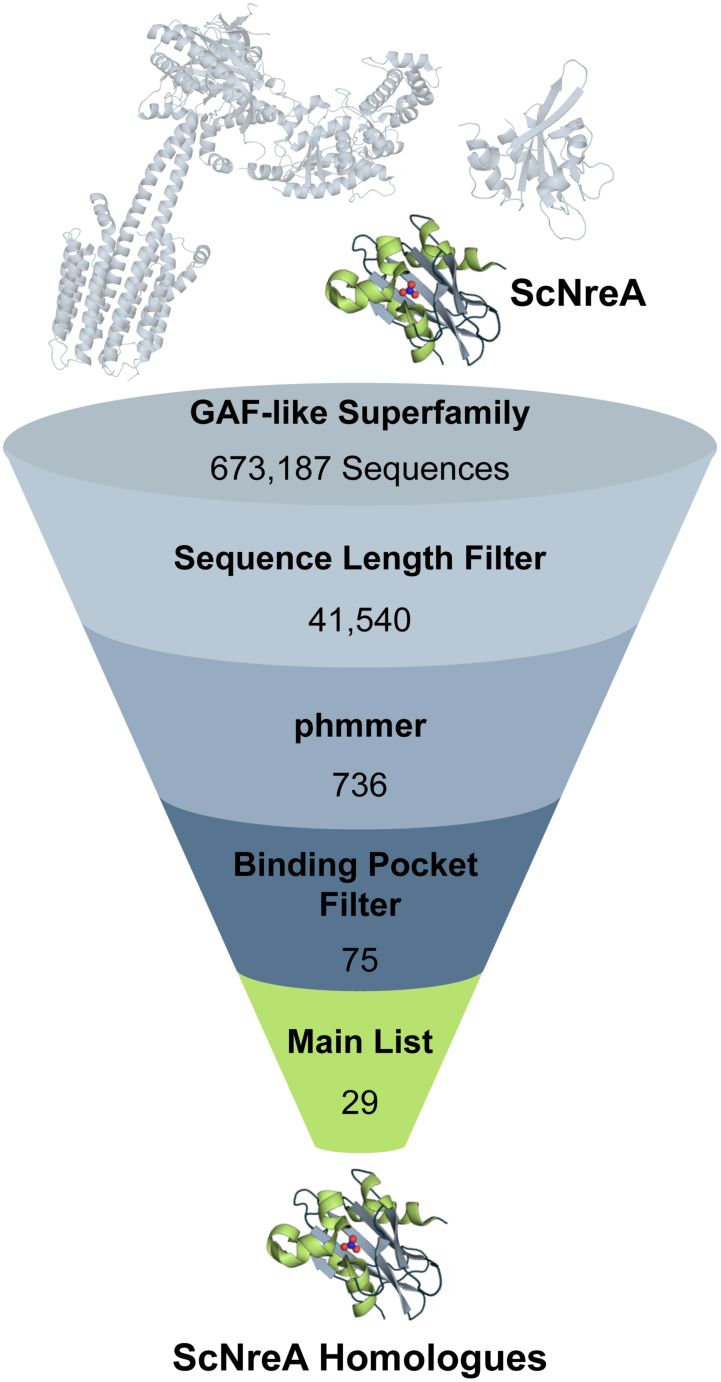
Bioinformatic pipeline used to identify ScNreA homologues within the GAF-like superfamily (InterPro, IPR029216). Sequences were filtered by length, analyzed with phmmer, and screened for conserved binding pocket residues W45, L67, A68, Y95, and I97 in ScNreA.

To examine how sequence variation influences nitrate recognition within a conserved biological context, we restricted our selection to staphylococcal homologues. Removal of all non-staphylococcal sequences, followed by elimination of redundant sequences from the same species, yielded 29 unique sequences. Analysis of the genomic context revealed that these homologues were commonly associated with genes encoding the NreB histidine kinase, NreC response regulator, NarGHJI nitrate reductase, and/or NarT transporter, supporting their roles in nitrate sensing (Figure S3) (Fedtke et al., 2002). From this set, five homologues were selected for thermodynamic characterization by ITC based on theoretical pI and percent sequence identity relative to ScNreA and SaNreA (Table S1). The selected homologues included NreAs from *S. simulans* (SsimNreA), *S. haemolyticus* (ShNreA), *S. argenteus* (SarNreA), *S. schleiferi* (SsNreA), and *S. aureus* (SaNreA), spanning 46–77.5% sequence identity to ScNreA. AlphaFold3 models of all five proteins were consistent with conservation of the overall GAF-like fold and nitrate-binding pocket, with coordinating residues adopting similar orientations to those in ScNreA, supporting the selection of these homologues for experimental characterization (Figure S5, S6).

### Nitrate-binding Thermodynamics of NreA Homologues

Each homologue was expressed in *Escherichia coli* with a C-terminal polyhistidine tag and purified by affinity and size-exclusion chromatography, yielding monomeric proteins (Figure S15–S19). Nitrate-binding thermodynamics were determined for all five proteins by isothermal titration calorimetry at 20 °C in 50 mM HEPES buffer (pH 7.8) containing 50 mM NaCl (Table 1; Figure S21–S26; Table S5–S10). These data were compared with those previously reported for ScNreA by our group. For all five homologues, the data were well fit by a 1:1 binding model, with favorable free energy changes (ΔG). Based on the ΔG and corresponding nitrate-binding affinities, the six proteins resolved into two distinct groups. ScNreA, SsimNreA, and ShNreA had the most favorable ΔG (–31.3 to –29.9 kJ/mol), corresponding to strong, low-micromolar binding affinities (*K*_d_ = 2.7 to 4.8 µM) (Ji et al., 2025). Conversely, SarNreA, SsNreA, and SaNreA had less favorable ΔG (–24.0 to –22.4 kJ/mol) and weaker, higher-micromolar binding affinities (*K*_d_ = 53.1 to 101.5 µM).

**Table 1.** Thermodynamic parameters of nitrate binding to ScNreA and selected homologues at 20 °C in 50 mM HEPES buffer (pH 7.8) with 50 mM NaCl.*^a^*.

| Protein | Sequence Identity to ScNreA (%) | $K_d$ (μM) | $\Delta H$ (kJ/mol) | $T\Delta S$ (kJ/mol) | $\Delta G$ (kJ/mol) |
| --- | --- | --- | --- | --- | --- |
| ScNreA<sup>b</sup> | – | $2.7 \pm 0.1$ | $-30.1 \pm 0.1$ | $1.3 \pm 0.1$ | $-31.3 \pm 0.1$ |
| SsimNreA | 77.5 | $4.8 \pm 0.1$ | $-26.4 \pm 0.1$ | $3.6 \pm 0.1$ | $-29.9 \pm 0.1$ |
| ShNreA | 51.0 | $4.7 \pm 0.6$ | $-24.4 \pm 1.2$ | $5.6 \pm 1.6$ | $-30.0 \pm 0.4$ |
| SarNreA | 47.3 | $94.6 \pm 12.1$ | $-11.7 \pm 0.4$ | $11.0 \pm 0.6$ | $-22.6 \pm 0.3$ |
| SsNreA | 46.0 | $101.5 \pm 6.1$ | $-25.6 \pm 2.2$ | $-3.2 \pm 2.1$ | $-22.4 \pm 0.1$ |
| SaNreA | 46.0 | $53.1 \pm 2.5$ | $-21.3 \pm 2.6$ | $2.8 \pm 2.5$ | $-24.0 \pm 0.1$ |

<sup>a</sup>The data are reported as the average ± standard deviation from two protein preparations, each determined by analysis of three technical replicates.
<sup>b</sup>ScNreA data were previously reported (Ji et al., 2025).

Further analysis of the individual thermodynamic parameters revealed that nitrate binding was exothermic for all six proteins, as indicated by negative enthalpy changes (ΔH), with distinct entropic terms (TΔS). The ΔH ranged from –30.1 to –11.7 kJ/mol, with ScNreA and SarNreA representing the extremes (Ji et al., 2025). For the high-affinity group, the ΔH became progressively less favorable from ScNreA to SsimNreA to ShNreA and was offset by increasingly favorable TΔS, resulting in comparable ΔG. In contrast, no clear trend in the balance between ΔH and TΔS was observed across the low-affinity group. SaNreA had a favorable TΔS value within the range measured for the high-affinity group, but a less favorable ΔH. While SsNreA retained a comparatively favorable ΔH, it had an unfavorable TΔS. Finally, SarNreA had the least favorable ΔH despite having the most favorable TΔS.

To better understand the basis for the distinct affinity groups, we determined the heat-capacity changes (ΔC_p_) associated with nitrate binding. ScNreA and SaNreA were selected as representative high- and low-affinity homologues, respectively. Temperature-dependent ITC measurements for SaNreA at 10, 15, 20, and 25 °C were used to determine ΔC_p_, which was compared with our previously reported value for ScNreA (Figure S25, S26; Table S9, S10). Upon nitrate binding, SaNreA exhibited a more negative ΔC_p_ (–733.5 J/mol•K) than ScNreA (–308.2 J/mol•K). This difference prompted us to consider structural features that might contribute to their distinct thermodynamic responses. Given the previously reported conformational variability of the SaNreA C-terminal region, we re-examined the nitrate-bound X-ray crystal structures of ScNreA and SaNreA and identified differences in tertiary contacts between the C-terminal region and the protein core (Figure S31).

### C-terminal Chimeragenesis Alters Nitrate-Binding Affinity

To determine whether C-terminal differences extended across the homologues, we generated AlphaFold3 models (Figure S32). ScNreA was predicted to have a loop-like C-terminus whereas SaNreA was predicted to have an α-helical C-terminus, like the Y94A apo-like mutant. The C-termini were also predicted to adopt loop-like conformations for the high-affinity homologues SsimNreA and ShNreA and α-helical conformations for the low-affinity homologues SsNreA and SarNreA. Confidence scores in the predicted C-terminal regions varied among the models (Figure S32). Based on these observations, we hypothesized that the 16-residue C-terminal region could tune nitrate-binding affinity. To test this hypothesis, we designed reciprocal C-terminal chimeras between the high-affinity ScNreA and low-affinity SaNreA. Specifically, residues I140 to P155 in ScNreA were replaced with residues D135 to K150 from SaNreA to generate the ScNreA chimera whereas residues D135 to K150 in SaNreA were replaced with residues I140 to P155 from ScNreA to generate the SaNreA chimera (Figure 3; Figure S13, S14). AlphaFold3 models of the ScNreA chimera predicted an α-helical C-terminus versus a loop-like C-terminus for the SaNreA chimera (Figure S33).

**Figure 3.**
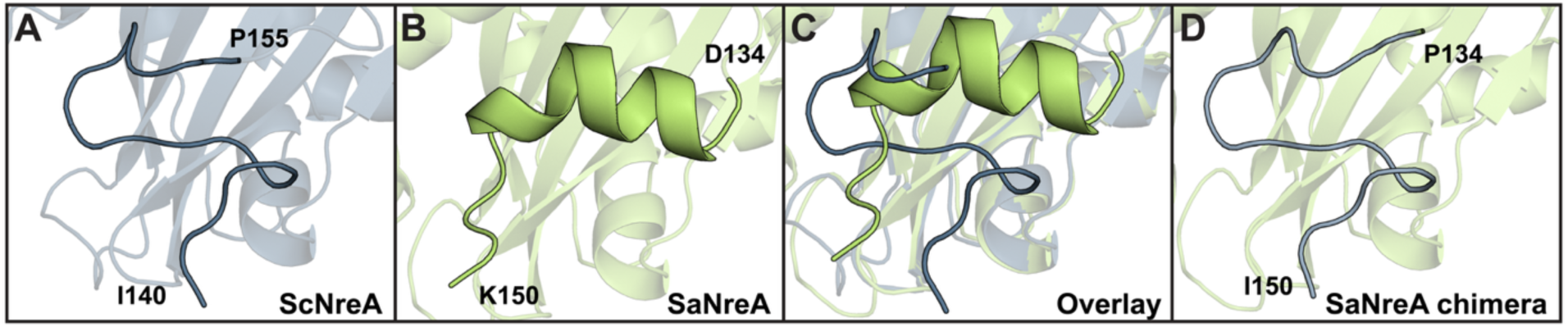
AlphaFold3 models of (A) ScNreA and (B) SaNreA highlighting predicted differences in the secondary structure of the C-terminal region. (C) Overlay of the ScNreA and SaNreA models shown in panels A and B. (D) AlphaFold3 model of the SaNreA chimera with the C-terminal region from ScNreA. Residue identities and sequence positions marking the start and end of the C-terminal regions are indicated.

Both chimeras were expressed, but only the SaNreA chimera could be isolated for characterization (Figure S20). The ScNreA chimera was therefore not further pursued. Far-ultraviolet circular dichroism spectra of SaNreA and the SaNreA chimera showed similar overall spectral profiles in the absence and presence of nitrate, indicating that the C-terminal replacement did not dramatically alter the overall secondary structure of the protein (Figure S29, S30). At 20 °C, the chimera had an intermediate ΔG (–27.9 kJ/mol) between SaNreA (–24.0 kJ/mol) and ScNreA (–31.3 kJ/mol), with an approximately five-fold increase in affinity (*K*_d_ = 10.4 µM) relative to SaNreA (Table 2; Figure S27, S28; Table S11, S12) (Ji et al., 2025). Nitrate binding to the SaNreA chimera was exothermic, with a favorable ΔH that was partially offset by an unfavorable TΔS term. This thermodynamic relationship was distinct from those of both SaNreA and ScNreA. However, the ΔC_p_ of the SaNreA chimera (–601.7 J/mol•K) remained closer in magnitude to SaNreA than to ScNreA (Table 2).

**Table 2.**
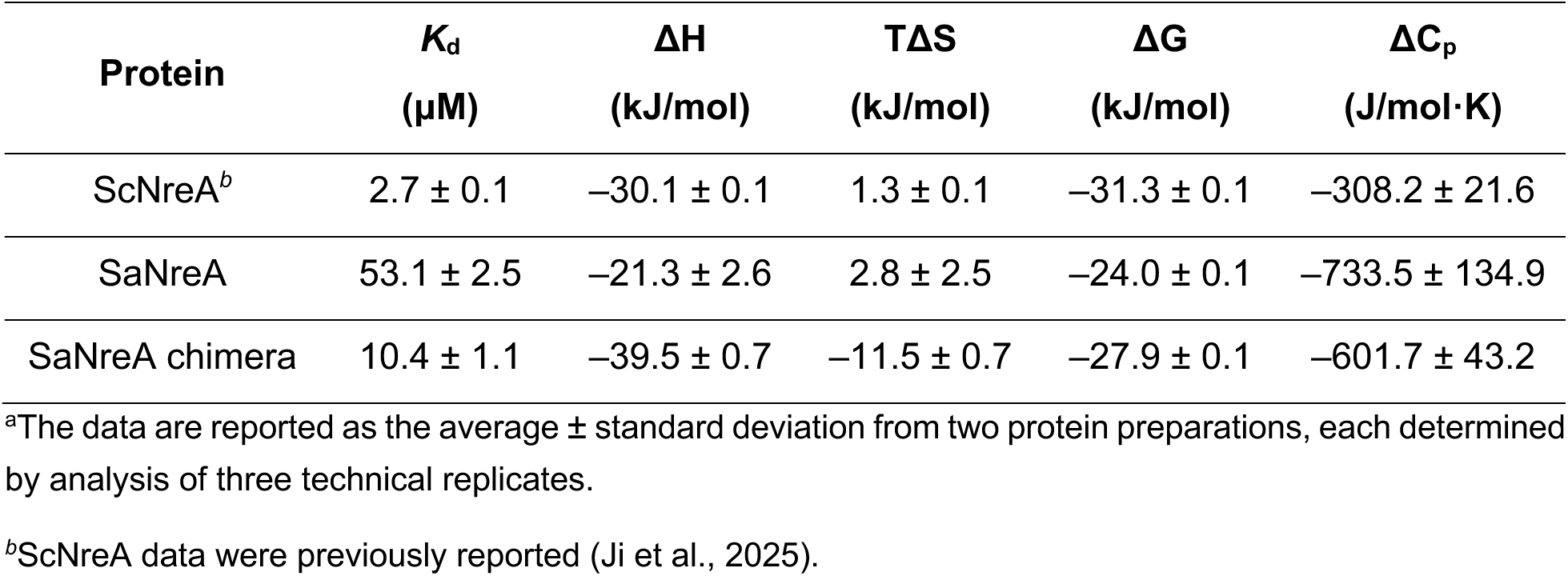
Thermodynamic parameters of nitrate binding to ScNreA, SaNreA, and the SaNreA chimera at 20 °C, with heat-capacity changes determined from temperature-dependent measurements, in 50 mM HEPES buffer (pH 7.8) with 50 mM NaCl.^a^.

## Discussion

In this study, we show that a conserved nitrate-binding pocket in staphylococcal NreA sensors supports nitrate binding with diverse thermodynamic profiles. Like ScNreA, all homologues bound nitrate exothermically. Interestingly, the NreA homologues separated into two affinity groups: a low-micromolar group comprising ScNreA, SsimNreA, and ShNreA and a high-micromolar group comprising SarNreA, SsNreA, and SaNreA. Across the six proteins, nitrate-binding affinities spanned approximately 38-fold, with *K*_d_ values ranging from 2.7 to 101.5 µM. Within the selected panel, overall sequence identity did not track with nitrate-binding affinity. Although SsimNreA shared a relatively high sequence identity with ScNreA (77.5%), the remaining homologues spanned a narrow range (46.0–51.0%). Moreover, clear differences emerged in the enthalpy-entropy balance across the homologues, suggesting distinct contributions from nitrate-protein interactions, conformational reorganization, and solvent rearrangement.

Comparison of the representative high- and low-affinity homologues showed that the magnitude of the heat-capacity change for SaNreA was more than twice that measured for ScNreA, consistent with greater changes in hydration and/or conformational organization in response to nitrate binding in SaNreA (Du et al., 2016; Prabhu & Sharp, 2005; Vega et al., 2016). Because the homologues are predicted to preserve the overall GAF-like fold and retain the same nitrate-binding pocket residues, these thermodynamic differences point to structural features outside the conserved binding pocket as potential contributors to nitrate-binding affinity.

Examination of the available X-ray crystal structures revealed differences in the conformation and tertiary interactions of the C-terminal region between ScNreA and SaNreA (Figure S1; Figure S31) (Niemann et al., 2014; Sangare et al., 2020). Extending this analysis across the homologues, AlphaFold3 models suggested a trend toward loop-like C-termini for the high-affinity NreAs and α-helical C-termini for the low-affinity NreAs (Figure S32). These observations led us to test the C-terminal region directly. Replacement of the 16-amino acid SaNreA C-terminal region (10.7% of the protein) with the corresponding region from ScNreA increased the nitrate-binding affinity five-fold and shifted the enthalpy-entropy balance, while the overall heat-capacity change remained closer to that of SaNreA than ScNreA. Together, these data provide direct experimental evidence that the non-coordinating C-terminal region can tune nitrate-binding thermodynamics.

Given this marked effect, we considered how changes at the sequence level could contribute. The three low-affinity homologues share highly similar C-terminal sequences, consistent with their sequence similarity to one another, whereas the high-affinity homologues do not share an obvious C-terminal consensus (Table S15). Moreover, no clear physicochemical pattern emerged across the C-terminal sequences that distinguished the two groups. Rather, differences in the C-terminal sequence and predicted conformation may influence how this region packs against and interacts with the surrounding protein scaffold (Figure S31) (Sangare et al., 2020). Thus, the C-terminus does not appear to encode affinity in a simple manner but instead represents an evolutionarily flexible element.

This is particularly relevant to the sensor function of NreA in staphylococci, where nitrate binding modulates the interaction between NreA and the NreB histidine kinase (Nilkens et al., 2014). Given that the C-terminal region of SaNreA has been proposed to contribute to this interaction, our findings raise the possibility that variation in nitrate-binding affinity tunes the nitrate concentration range over which NreA relieves inhibition of NreB, thereby promoting nitrate-responsive gene expression (Sangare et al., 2020). Investigation of matched ligand-free and nitrate-bound states across additional homologues, together with direct measurements of NreA-NreB interactions and downstream signaling, will help define how the C-terminus couples nitrate sensing to regulatory function.

While our investigation focused on staphylococcal NreAs, we identified over 700 ScNreA homologues across diverse bacterial taxa, spanning 15.6–99.4% sequence identity relative to ScNreA. Inspection of representative genome neighborhoods further showed that these homologues occur not only alongside genes encoding the established staphylococcal NreB/NreC two-component regulatory system, NarT transporter, and NarGHJI reductase but also within diverse regulatory, transport, and metabolic contexts (Figure S34). These observations suggest that NreA-like nitrate sensing may extend across a broader range of bacterial systems than has been experimentally characterized and highlight unexplored functional contexts across this protein family.

## Data Availability

The data that support the findings of this study are available in the Main Text and Supporting Information of this article. The corresponding author can be contacted for additional requests.

## Supporting Information

Experimental data (PDF). The sequence FASTA files are available in a Zenodo repository (DOI 10.5281/zenodo.22907091).

## Author Contributions

E.K.P.: data curation, formal analysis, investigation, methodology, validation, visualization, writing – original draft, writing – review & editing. K.J.: data curation, methodology, supervision, writing – review & editing. J.M.: methodology. W.P.: data curation, methodology. M.A.C.: data curation, methodology. G.M.: methodology, writing – review & editing. S.C.D.: conceptualization, funding acquisition, supervision, writing – review & editing.

## Acknowledgements

The authors acknowledge Dr. Shelby M. Phelps for assistance with molecular cloning of the SaNreA chimera and Mr. Dylan M. Wicherts for editorial feedback. S.C.D. acknowledges the National Institute of General Medical Sciences (R35GM128923) and the Welch Foundation (AT-1918–20170325 and AT-2060-20210327) for support of preliminary studies and personnel (K.J., E.K.P., W.P., and M.A.C.) that established the foundation for this work; the Welch Foundation (AT-2060-20240404) for support of personnel (K.J.); the National Science Foundation (2240095) for support of personnel (E.K.P.) and primary support of the research reported here. J.M. acknowledges the National Institute of Allergy and Infectious Diseases (R01AI178692) for personnel support. This study does not represent the views of the supporting agency and is the responsibility of the authors.

## AI Disclosure

ChatGPT was used for general editing, and the output was reviewed by the authors.

