## Supporting Information for "Thermodynamic Diversity in Staphylococcal Nitrate Sensors with a Conserved Binding Pocket"

### X-Ray Crystal Structures

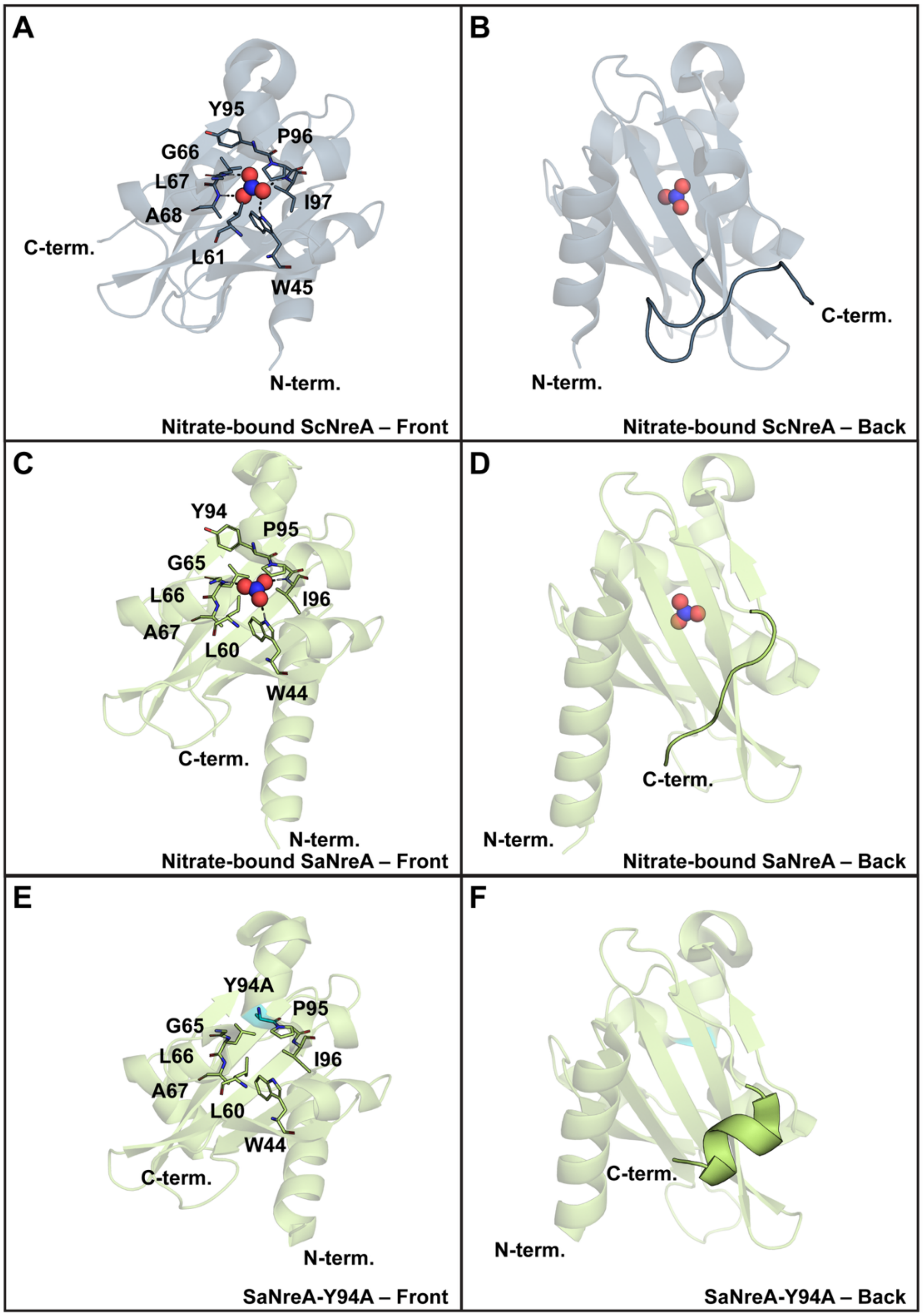

**Figure S1.** Front (left) and back views (right) of the crystal structures of (A–B) nitrate-bound ScNreA (PDB ID: 4IUK), (C–D) nitrate-bound SaNreA (PDB ID: 6IZJ), and (E–F) SaNreA-Y94A (PDB ID: 6K2H). The front views highlight residues corresponding to those within 4 Å of the nitrate in the ScNreA crystal structure, and the back views highlight the differences in the secondary structure of the C-terminal region. The 1,2-ethanediol (EDO) molecules are omitted for clarity from the SaNreA and SaNreA-Y94A crystal structures.

### C-Terminal Electrostatic Maps and Key Residues

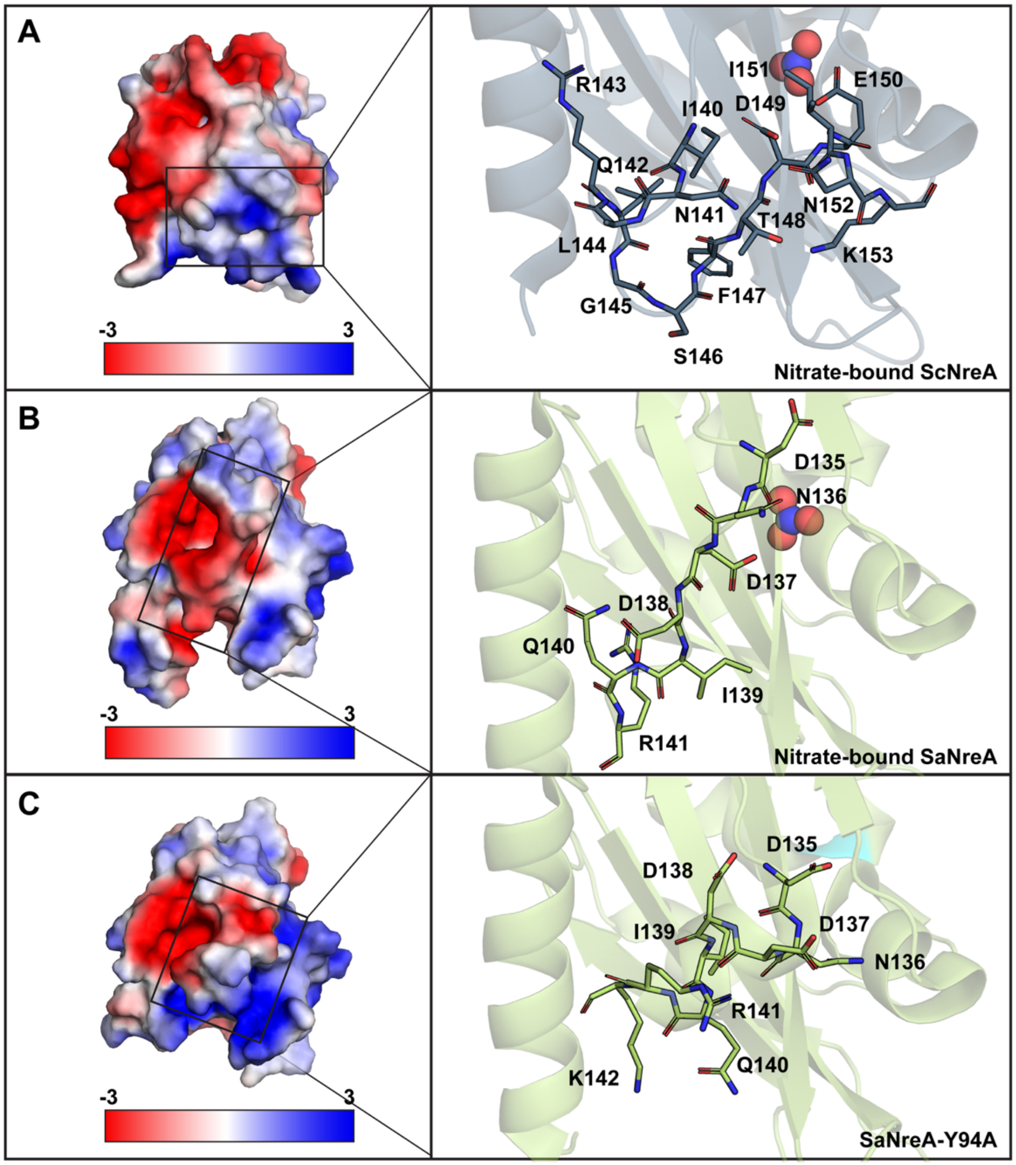

**Figure S2.** Comparison of the C-terminal electrostatic surface potentials (left) with insets (right) highlighting key residues as sticks for (A) nitrate-bound ScNreA (PDB ID: 4IUK), (B) nitrate-bound SaNreA (PDB ID: 6IZJ), and (C) SaNreA-Y94A (PDB ID: 6K2H). The EDO molecules are omitted for clarity from the SaNreA and SaNreA-Y94A crystal structures.

### Table of NreA Homologues from Bioinformatic Search

**Table S1.** ScNreA homologues identified through bioinformatic mining that retain the ScNreA nitrate-binding pocket residues with their percent sequence identity to ScNreA and SaNreA and theoretical isoelectric points. The homologues selected for experimental testing are highlighted in green.

| UniProt Accession Number | Organism | Percent Identity Compared to ScNreA | Percent Identity Compared to SaNreA | Theoretical Isoelectric Point |
|---|---|---|---|---|
| B9DL91 | *S. carnosus* (Strain TM300) | – | 46.0 | 6.7 |
| A0A143PE72 | *S. condimenti* | 92.8 | 48.0 | 8.0 |
| A0A239TS44 | *S. piscifermentans* | 84.9 | 49.3 | 7.8 |
| A0A855LGD0 | *S. simulans* | 77.5 | 50.0 | 6.2 |
| A0A5B2Z1J8 | *Staphylococcus sp.* GDX7P459A | 56.2 | 59.1 | 6.5 |
| A0A1D4M2Y6 | *S. caeli* | 55.0 | 54.0 | 6.9 |
| A0A1E9A7V5 | *Staphylococcus sp.* HMSC071G07 | 54.4 | 53.3 | 7.2 |
| A0A1Z3U0P9 | *S. pettenkoferi* | 54.4 | 53.3 | 7.2 |
| A0A2K4FC37 | *S. argensis* | 54.4 | 53.3 | 7.2 |
| A0A2T4LSE0 | *S. cohnii* | 53.3 | 56.7 | 9.1 |
| A0A2T4SB76 | *S. nepalensis* | 53.3 | 54.7 | 9.3 |
| A0A418IQ79 | *S. xylosus* | 51.3 | 58.7 | 9.0 |
| Q4L8Q8 | *S. haemolyticus* (Strain JCSC1435) | 51.0 | 58.4 | 7.1 |
| A0A418ID80 | *S. shinii* | 50.7 | 58.7 | 9.2 |
| A0A418HR02 | *S. gallinarum* | 50.0 | 56.7 | 6.4 |
| A0A2K4DTF5 | *S. devriesei* | 49.5 | 58.4 | 6.7 |
| A0A654BWP9 | *Staphylococcus sp.* 8AQ | 49.5 | 61.3 | 8.6 |
| A0A0M2NLG6 | *S. pasteuri* | 49.5 | 61.3 | 8.5 |
| A0A2C6WML9 | *S. edaphicus* | 49.3 | 54.7 | 9.2 |
| A0A380FW79 | *S. petrasii* | 48.5 | 56.4 | 6.4 |
| K9AZ44 | *S. massiliensis* S46 | 47.7 | 45.0 | 6.7 |
| G5JKI0 | *S. simiae* (CCM 7213 = CCUG 51256) | 47.7 | 85.2 | 8.0 |
| A0A7U7PWK7 | *S. argenteus* | 47.3 | 94.0 | 8.7 |
| A0A4Q9WQ72 | *S. hominis* | 46.7 | 57.3 | 9.4 |
| A0A7Z7W3U1 | *S. schleiferi* | 46.0 | 98.7 | 9.4 |
| A0A2K4AG13 | *S. schweitzeri* | 46.0 | 99.3 | 9.3 |
| Q2FVM5 | *S. aureus* (Strain NCTC 8325 / PS 47) | 46.0 | – | 9.4 |
| A0A292DHU4 | *S. lugdunensis* | 45.0 | 54.4 | 7.9 |
| A0A4Z1BY69 | *S. pragensis* | 42.3 | 58.4 | 7.1 |

### Genome Neighborhood Diagrams for NreA Homologues

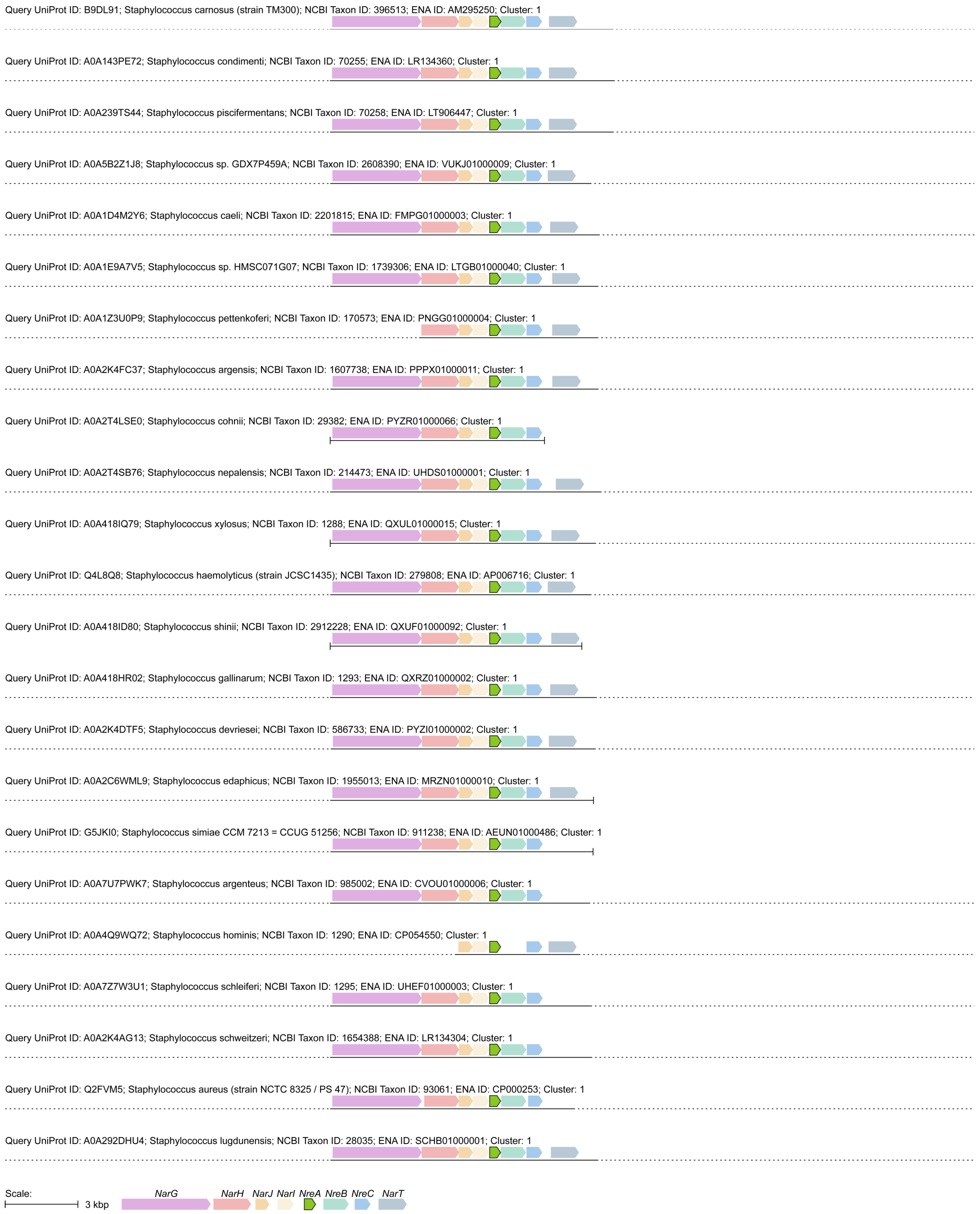

**Figure S3.** Genome neighborhood diagrams for the putative NreA homologues shown in Table S1. For each diagram, the *nreA* gene (green) is highlighted along with neighboring genes encoding the NreB histidine kinase (*nreB*, mint), NreC response regulator (*nreC*, blue), NarGHJI nitrate reductase (*narG*, purple; *narH*, pink; *narJ*, orange; *narI*, yellow), and NarT nitrate transporter (*narT*, gray-blue). Homologues from Table S1 no longer found in UniProt were excluded.

### Multiple Sequence Alignment for Selected NreAs and the SaNreA Chimera

```
ScNreA              MLNSVIASDYFDYQDALDEIRETEKFDFAAIALPEDGLHSAVIKWKYASGNINYRYRMIV      60
SsimNreA            -MNSVVATGYFNYQVALEEIREEEQFDFAAIALPEDDSHSAVIKWRHASGNLNNRYQLIV      59
ShNreA              -MITNPVFEKQDFQQQLNNLRVSEGYDFAGVALYEHHMTSAPIKWHYVSGNLNDRYKMII      59
SarNreA             -MTPEAMIEDQSFQETLDKIRKEEGYDFAAIAFYESNKPSSPIKWHYVSGNKNNRFKLII      59
SsNreA              -MTPEAIIEDRRFQETLDKIRKEEGYDFAAIAFYESNKPSSPIKWHYVSGNKNNRFKLII      59
SaNreA              -MTPEAIIEDRRFQETLDKIRKEEGYDFAAIAFYESNKPSSPIKWHYVSGNKNNRFKLII      59
SaNreA_chimera      -MTPEAIIEDRRFQETLDKIRKEEGYDFAAIAFYESNKPSSPIKWHYVSGNKNNRFKLII      59
                     :          :*  *:::*  * :***.:*: *    *: ***::.*** * *:::*:

ScNreA              LRPGKGLAGLVIRTGSRKIVEDVDAELSQNDKLGYPIVLSEALTAMVAIPLWKNNRVYGA      120
SsimNreA            LRPGKGLAGLVIRTGSRMIIDDVNTVLSPNDKLSYPIVLSESLTAMVAIPLWHDNRVYGA      119
ShNreA              LRKGRGLAGMVMKTGKRMIIADVVASLSPEDKIKYPILIAENLTAMVAIPLCYNNQVYGV      119
SarNreA             LRKGKGLAGTVMKTGKRMIIANVGLALGPEEKIDYPILLSESLTAVFAVPLWYKNQVYGV      119
SsNreA              LRKGRGLAGTVMKTGKRMIIANVGLALGPEEKIDYPILLSESLTAVLAVPLWYKNQVYGV      119
SaNreA              LRKGRGLAGTVMKTGKRMVIANVGLALGPEEKIDYPILLSESLTAVLAVPLWYKNQVYGV      119
SaNreA_chimera      LRKGRGLAGTVMKTGKRMVIANVGLALGPEEKIDYPILLSESLTAVLAVPLWYKNQVYGV      119
                    ** *:**** *::**.* :: :*   *. ::*: ***:::* ***:.*:**  .*:***.

ScNreA              LLLGQREGRPLPEGSTTFRINQRLGSFTDEINKQP  155
SsimNreA            LLLGQRNGRPLPEGSEEIRINHRLGSFRDEIN---  151
ShNreA              LLLGQRDGKSLPDYETL-DISGKLWTFSEEM----  149
SarNreA             LLFGQRDGRPLPKIFDYEDIQYKFGIFNDDK----  150
SsNreA              LLFGQRDGRSLPKIFDNDDIQRKFGIFNDDK----  150
SaNreA              LLFGQRDGRPLPKIFDNDDIQRKFGIFNDDK----  150
SaNreA_chimera      LLFGQRDGRPLPKIFINQR----LGSFTDEINKQP  150
                    **:***:*: **.          :  * ::
```

**Figure S4.** Multiple sequence alignment of the selected NreAs and the SaNreA chimera characterized in this study. Residues corresponding to those within 4 Å of nitrate in the ScNreA crystal structure are highlighted in green, and the 16 amino acids defined as the C-terminal region in this study are highlighted in yellow.

### Front Views of the AlphaFold3 Models with Confidence Levels

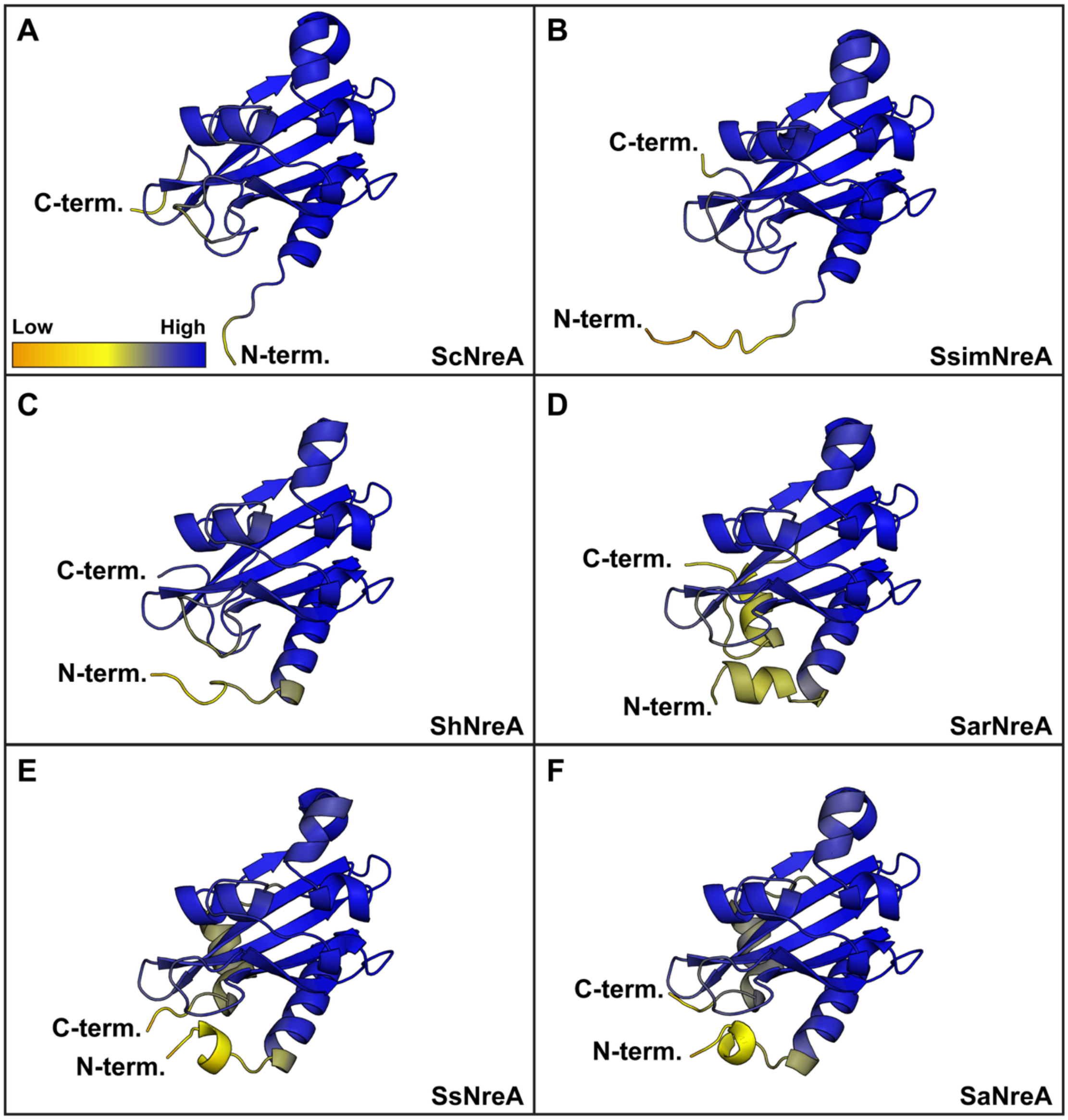

**Figure S5.** Front views of the AlphaFold3 models highlighting the predicted model confidence for (A) ScNreA, (B) SsimNreA, (C) ShNreA, (D) SarNreA, (E) SsNreA, and (F) SaNreA with dark blue representing high confidence, and bright orange representing low confidence. Residues M1–I6 of ScNreA are not shown for clarity at the displayed figure scale.

### AlphaFold3 Models Highlighting the Nitrate-Binding Pockets

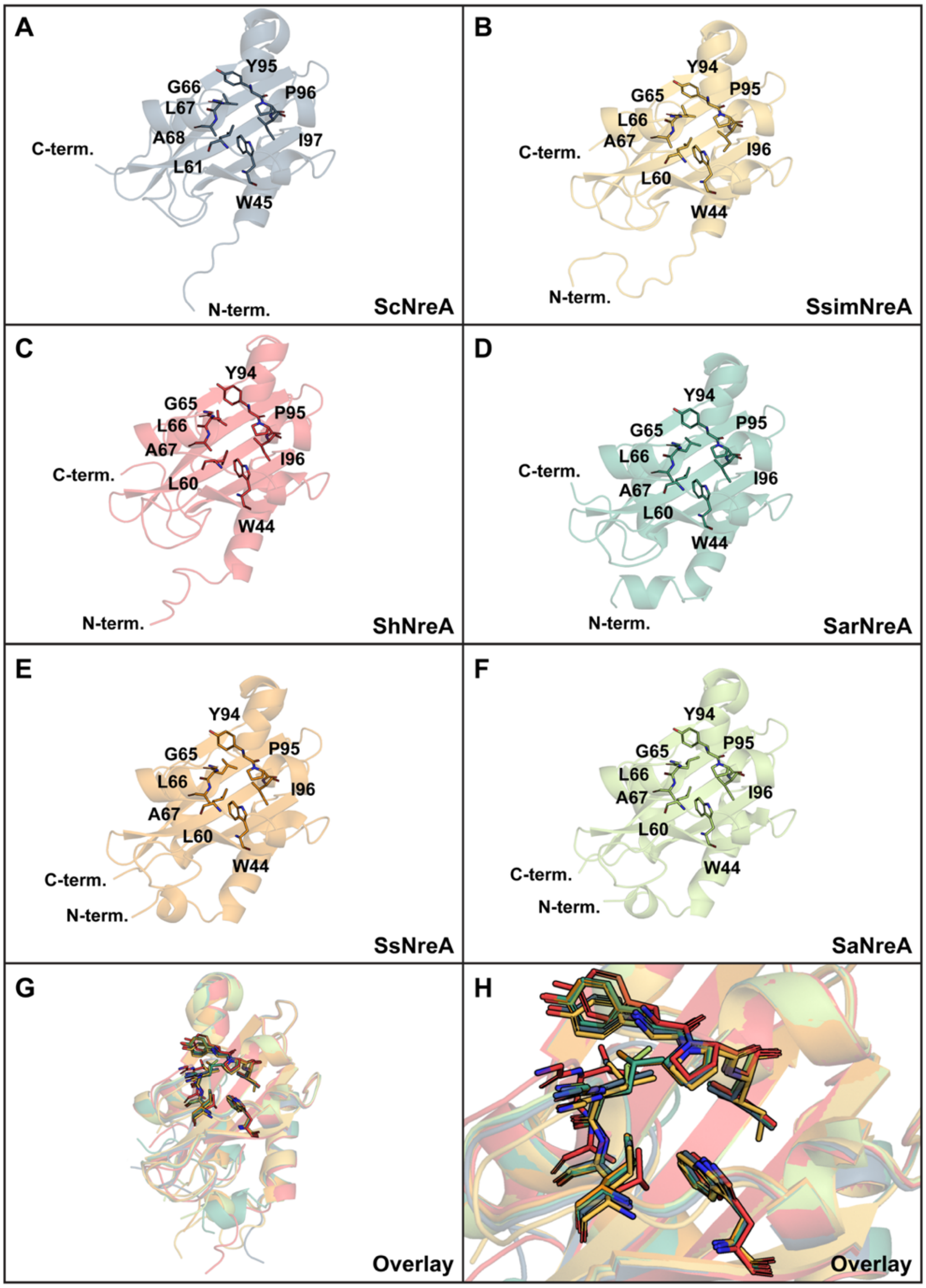

**Figure S6.** AlphaFold3 models of (A) ScNreA (blue), (B) SsimNreA (yellow), (C) ShNreA (red), (D) SarNreA (turquoise), (E) SsNreA (orange), and (F) SaNreA (green), showing residues corresponding to those within 4 Å of nitrate in the ScNreA crystal structure. Overlay of the models highlighting the (G) conserved GAF-like fold and (H) nitrate-binding pockets. Residues M1–I6 of ScNreA are not shown for clarity at the displayed figure scale.

### Nucleotide and Amino Acid Sequences

**CATATG**CTGAACAGCGTGATCGCGAGCGACTACTTCGATTATCAGGACGCGCTGG
ATGAGATCCGTGAGACCGAAAAGTTCGACTTTGCGGCGATTGCGCTGCCGGAAGA
TGGTCTGCACAGCGCGGTTATTAAGTGGAAATACGCGAGCGGCAACATCAACTAC
CGTTATCGTATGATTGTGCTGCGTCCGGGCAAGGGTCTGGCGGGTCTGGTTATCC
GTACCGGCAGCCGTAAAATTGTGGAGGACGTTGATGCGGAACTGAGCCAGAACGA
CAAGCTGGGTTATCCGATTGTGCTGAGCGAGGCGCTGACCGCGATGGTTGCGATT
CCGCTGTGGAAAAACAACCGTGTGTATGGTGCGCTGCTGCTGGGTCAACGTGAAG
GTCGTCCGCTGCCGGAAGGTAGCACCACCTTCCGTATCAACCAGCGTCTGGGCAG
CTTTACCGATGAAATTAACAAACAACCG**AAGCTT**GCGGCCGCACTCGAG**CACCAC**
**CACCACCACCAC****TGA**

| | | |
|---|---|---|
| 1 | MLNSVIASDYFDYQDALDEI | 20 |
| 21 | RETEKFDFAAIALPEDGLHS | 40 |
| 41 | AVIKWKYASGNINYRYRMIV | 60 |
| 61 | LRPGKGLAGLVIRTGSRKIV | 80 |
| 81 | EDVDAELSQNDKLGYPIVLS | 100 |
| 101 | EALTAMVAIPLWKNNRVYGA | 120 |
| 121 | LLLGQREGRPLPEGSTTFRI | 140 |
| 141 | NQRLGSFTDEINKQPKLAAA | 160 |
| 161 | LE**HHHHHH*** | 168 |

**Figure S7.** Nucleotide sequence (top) and translated amino acid sequence (bottom) of ScNreA used in this study. Vector-derived restriction sites (black, bold), the polyhistidine tag (purple, bold), and the stop codon (red, bold) are indicated.

**CATATG**AATTCAGTTGTAGCTACAGGATATTTTAACTATCAGGTTGCGTTAGAAGAG
ATCAGAGAAGAGGAGCAATTTGACTTCGCCGCAATCGCGCTGCCGGAAGATGACA
GCCACAGCGCAGTCATCAAGTGGCGTCACGCTAGCGGTAATCTGAATAACCGCTA
CCAACTGATTGTTCTGCGTCCGGGTAAAGGCCTGGCCGGTTTGGTTATTCGTACG
GGTTCCCGGATGATTATCGACGACGTGAATACCGTGCTCTCTCCGAATGATAAACT
GTCCTACCCGATTGTGCTTAGTGAGAGCCTGACCGCTATGGTGGCGATTCCGCTG
TGGCATGATAACCGTGTTTATGGTGCGCTGTTGTTGGGCCAGCGTAACGGCCGTC
CACTGCCGGAAGGCAGCGAGGAGATCCGCATCAACCATCGTCTGGGCTCGTTCC
GCGATGAAATCAAC**AAGCTT**GCGGCCGCACTCGAG**CACCACCACCACCACCAC****T
GA**

| | | |
|---|---|---|
| 1 | MNSVVATGYFNYQVALEEIR | 20 |
| 21 | EEEQFDFAAIALPEDDSHSA | 40 |
| 41 | VIKWRHASGNLNNRYQLIVL | 60 |
| 61 | RPGKGLAGLVIRTGSRMIID | 80 |
| 81 | DVNTVLSPNDKLSYPIVLSE | 100 |
| 101 | SLTAMVAIPLWHDNRVYGAL | 120 |
| 121 | LLGQRNGRPLPEGSEEIRIN | 140 |
| 141 | HRLGSFRDEINKLAAALE**HH** | 160 |
| 161 | **HHHH*** | 164 |

**Figure S8.** Nucleotide sequence (top) and translated amino acid sequence (bottom) of SsimNreA used in this study. Vector-derived restriction sites (black, bold), the polyhistidine tag (purple, bold), and the stop codon (red, bold) are indicated.

**CATATG**ATCACCAACCCGGTGTTCGAGAAGCAAGACTTTCAGCAACAGCTGAACAA
CCTGCGTGTGAGCGAGGGTTACGATTTCGCGGGCGTTGCGCTGTATGAACACCAC
ATGACCAGCGCGCCGATTAAATGGCACTACGTGAGCGGTAACCTGAACGACCGTT
ATAAGATGATCATTCTGCGTAAAGGTCGTGGCCTGGCGGGTATGGTTATGAAGAC
CGGCAAACGTATGATCATTGCGGATGTGGTTGCGAGCCTGAGCCCGGAGGATAAG
ATCAAATACCCGATCCTGATTGCGGAAAACCTGACCGCGATGGTTGCGATTCCGCT
GTGCTACAACAACCAAGTGTATGGTGTTCTGCTGCTGGGTCAGCGTGATGGCAAG
AGCCTGCCGGACTATGAAACCCTGGACATCAGCGGCAAACTGTGGACCTTTAGCG
AGGAAATG**AAGCTT**GCGGCCGCACTCGAG**CACCACCACCACCACCAC****TGA**

| | | |
|---|---|---|
| 1 | MITNPVFEKQDFQQQLNNLR | 20 |
| 21 | VSEGYDFAGVALYEHHMTSA | 40 |
| 41 | PIKWHYVSGNLNDRYKMIIL | 60 |
| 61 | RKGRGLAGMVMKTGKRMIIA | 80 |
| 81 | DVVASLSPEDKIKYPILIAE | 100 |
| 101 | NLTAMVAIPLCYNNQVYGVL | 120 |
| 121 | LLGQRDGKSLPDYETLDISG | 140 |
| 141 | KLWTFSEEMKLAAALE**HHHH** | 160 |
| 161 | **HH*** | 162 |

**Figure S9.** Nucleotide sequence (top) and translated amino acid sequence (bottom) of ShNreA used in this study. Vector-derived restriction sites (black, bold), the polyhistidine tag (purple, bold), and the stop codon (red, bold) are indicated.

**CATATG**ACACCCGAAGCTATGATAGAGGATCAATCTTTTCAAGAAACGCTCGATAA
GATCCGTAAGGAGGAAGGTTACGACTTCGCGGCAATTGCCTTTTATGAGAGCAATA
AACCGAGCTCGCCGATTAAATGGCACTACGTGAGCGGTAATAAGAACAACCGCTTT
AAACTGATCATCCTGCGTAAGGGCAAGGGCTTGGCGGGCACCGTTATGAAGACCG
GTAAACGTATGATTATCGCTAATGTCGGTCTGGCGCTGGGTCCGGAAGAGAAAATT
GATTACCCGATCTTGCTGAGCGAATCCTTAACCGCGGTGTTCGCTGTGCCGCTGT
GGTATAAGAACCAGGTTTATGGTGTTCTGTTGTTTGGCCAACGCGACGGCCGTCCA
CTGCCGAAAATCTTCGACTACGAGGATATTCAGTATAAATTCGGTATTTTCAACGAT
GACAAG**AAGCTT**GCGGCCGCACTCGAG**CACCACCACCACCACCAC****TGA**

| | | |
|---|---|---|
| 1 | MTPEAMIEDQSFQETLDKIR | 20 |
| 21 | KEEGYDFAAIAFYESNKPSS | 40 |
| 41 | PIKWHYVSGNKNNRFKLIIL | 60 |
| 61 | RKGKGLAGTVMKTGKRMIIA | 80 |
| 81 | NVGLALGPEEKIDYPILLSE | 100 |
| 101 | SLTAVFAVPLWYKNQVYGVL | 120 |
| 121 | LFGQRDGRPLPKIFDYEDIQ | 140 |
| 141 | YKFGIFNDDKKLAAALE**HHH** | 160 |
| 161 | **HHH*** | 163 |

**Figure S10.** Nucleotide sequence (top) and translated amino acid sequence (bottom) of SarNreA used in this study. Vector-derived restriction sites (black, bold), the polyhistidine tag (purple, bold), and the stop codon (red, bold) are indicated.

**CATATG**ACACCCGAGGCTATAATTGAAGATAGGAGATTCCAAGAAACCCTGGATAA
GATCCGCAAAGAAGAGGGCTATGATTTTGCGGCAATTGCGTTTTATGAGTCCAATA
AGCCGAGCAGCCCGATCAAATGGCACTACGTGAGCGGTAATAAGAACAACCGCTT
CAAACTGATTATTCTGCGTAAGGGCCGTGGTCTCGCGGGTACGGTTATGAAAACC
GGTAAACGTATGATTATCGCCAATGTGGGCCTGGCTCTGGGTCCGGAAGAGAAAA
TCGATTACCCGATTCTGTTAAGCGAGTCGTTGACCGCAGTCTTGGCGGTGCCGCT
GTGGTACAAAAACCAGGTTTATGGTGTTCTTCTGTTTGGCCAGCGTGATGGTCGTT
CTTTGCCAAAGATCTTCGACAACGACGACATCCAACGCAAGTTCGGCATCTTTAAC
GACGACAAG**AAGCTT**GCGGCCGCACTCGAG**CACCACCACCACCACCAC****TGA**

| | | |
|---|---|---|
| 1 | MTPEAIIEDRRFQETLDKIR | 20 |
| 21 | KEEGYDFAAIAFYESNKPSS | 40 |
| 41 | PIKWHYVSGNKNNRFKLIIL | 60 |
| 61 | RKGRGLAGTVMKTGKRMIIA | 80 |
| 81 | NVGLALGPEEKIDYPILLSE | 100 |
| 101 | SLTAVLAVPLWYKNQVYGVL | 120 |
| 121 | LFGQRDGRSLPKIFDNDDIQ | 140 |
| 141 | RKFGIFNDDKKLAAALE**HHH** | 160 |
| 161 | **HHH*** | 163 |

**Figure S11.** Nucleotide sequence (top) and translated amino acid sequence (bottom) of SsNreA used in this study. Vector-derived restriction sites (black, bold), the polyhistidine tag (purple, bold), and the stop codon (red, bold) are indicated.

**CATATG**ACCCCGGAGGCGATCATTGAAGACCGTCGTTTCCAGGAGACCCTGGATA
AGATTCGTAAAGAGGAAGGTTACGACTTCGCGGCGATCGCGTTTTATGAAAGCAAC
AAGCCGAGCAGCCCGATTAAATGGCACTACGTGAGCGGCAACAAGAACAACCGTT
TTAAACTGATCATTCTGCGTAAGGGTCGTGGCCTGGCGGGTACCGTTATGAAGAC
CGGCAAACGTATGGTGATTGCGAACGTTGGTCTGGCGCTGGGTCCGGAGGAAAAA
ATCGATTATCCGATTCTGCTGAGCGAGAGCCTGACCGCGGTGCTGGCGGTTCCGC
TGTGGTACAAGAACCAGGTGTATGGTGTTCTGCTGTTCGGTCAACGTGACGGCCG
TCCGCTGCCGAAAATCTTTGATAACGACGATATTCAGCGTAAGTTCGGCATCTTTA
ACGACGATAAA**AAGCTTCACCACCACCACCACCACTGA**

| | | |
|---|---|---|
| 1 | MTPEAIIEDRRFQETLDKIR | 20 |
| 21 | KEEGYDFAAIAFYESNKPSS | 40 |
| 41 | PIKWHYVSGNKNNRFKLIIL | 60 |
| 61 | RKGRGLAGTVMKTGKRMVIA | 80 |
| 81 | NVGLALGPEEKIDYPILLSE | 100 |
| 101 | SLTAVLAVPLWYKNQVYGVL | 120 |
| 121 | LFGQRDGRPLPKIFDNDDIQ | 140 |
| 141 | RKFGIFNDDKKL**HHHHHH*** | 158 |

**Figure S12.** Nucleotide sequence (top) and translated amino acid sequence (bottom) of SaNreA used in this study. Vector-derived restriction sites (black, bold), the polyhistidine tag (purple, bold), and the stop codon (red, bold) are indicated.

**CATATG**CTAAATTCAGTAATAGCTAGTGATTATTTCGACTATCAAGATGCTCTGGAT
GAGATCCGCGAAACCGAAAAATTCGATTTTGCGGCAATCGCCCTTCCGGAGGACG
GTCTGCACAGCGCTGTGATTAAGTGGAAATACGCGAGCGGTAATATTAACTACCGT
TACCGCATGATTGTGCTGCGTCCGGGTAAAGGTTTAGCAGGCCTGGTTATCCGTA
CCGGCAGCCGTAAGATCGTAGAGGATGTTGATGCGGAACTGTCTCAAAACGACAA
ACTGGGTTATCCGATTGTCTTGTCCGAGGCGTTGACCGCCATGGTTGCGATCCCG
CTGTGGAAGAACAACCGCGTGTATGGTGCGCTCTTGCTGGGTCAGCGTGAGGGC
CGTCCACTGCCGGAAGGCTCGACTACGTTTCGT**GATAATGACGACATCCAGAGAA**
**AGTTCGGCATTTTTAACGACGACAAG****AAGCTT**GCGGCCGCACTCGAG**CACCACC**
**ACCACCACCAC****TGA**

| | | |
|---|---|---|
| 1 | MLNSVIASDYFDYQDALDEI | 20 |
| 21 | RETEKFDFAAIALPEDGLHS | 40 |
| 41 | AVIKWKYASGNINYRYRMIV | 60 |
| 61 | LRPGKGLAGLVIRTGSRKIV | 80 |
| 81 | EDVDAELSQNDKLGYPIVLS | 100 |
| 101 | EALTAMVAIPLWKNNRVYGA | 120 |
| 121 | LLLGQREGRPLPEGSTTFR**D** | 140 |
| 141 | **NDDIQRKFGIFNDDK**KLAAA | 160 |
| 161 | LE**HHHHHH*** | 168 |

**Figure S13.** Nucleotide sequence (top) and translated amino acid sequence (bottom) of the ScNreA chimera used in this study. Vector-derived restriction sites (black, bold), exchanged C-terminal sequence (blue, bold), polyhistidine tag (purple, bold), and stop codon (red, bold) are indicated.

**CATATG**ACCCCGGAGGCGATCATTGAAGACCGTCGTTTCCAGGAGACCCTGGATA
AGATTCGTAAAGAGGAAGGTTACGACTTCGCGGCGATCGCGTTTTATGAAAGCAAC
AAGCCGAGCAGCCCGATTAAATGGCACTACGTGAGCGGCAACAAGAACAACCGTT
TTAAACTGATCATTCTGCGTAAGGGTCGTGGCCTGGCGGGTACCGTTATGAAGAC
CGGCAAACGTATGGTGATTGCGAACGTTGGTCTGGCGCTGGGTCCGGAGGAAAAA
ATCGATTATCCGATTCTGCTGAGCGAGAGCCTGACCGCGGTGCTGGCGGTTCCGC
TGTGGTACAAGAACCAGGTGTATGGTGTTCTGCTGTTCGGTCAACGTGACGGCCG
TCCGCTGCCGAAAATCTTT**ATCAACCAGCGTCTGGGCAGCTTTACCGATGAAATT**
**AACAAACAACCGAAGCTTCACCACCACCACCACCACTGA**

| | | |
|---|---|---|
| 1 | MTPEAIIEDRRFQETLDKIR | 20 |
| 21 | KEEGYDFAAIAFYESNKPSS | 40 |
| 41 | PIKWHYVSGNKNNRFKLIIL | 60 |
| 61 | RKGRGLAGTVMKTGKRMVIA | 80 |
| 81 | NVGLALGPEEKIDYPILLSE | 100 |
| 101 | SLTAVLAVPLWYKNQVYGVL | 120 |
| 121 | LFGQRDGRPLPKIF**INQRLG** | 140 |
| 141 | **SFTDEINKQP**KL**HHHHHH*** | 158 |

**Figure S14.** Nucleotide sequence (top) and translated amino acid sequence (bottom) of the SaNreA chimera used in this study. Vector-derived restriction sites (black, bold), exchanged C-terminal sequence (blue, bold), polyhistidine tag (purple, bold), and stop codon (red, bold) are indicated.

### Table of PCR Primers

**Table S2.** Polymerase chain reaction (PCR) primers used to generate the SaNreA chimera plasmid.

| Description | Primer Sequence (5' → 3') |
|---|---|
| Insert Forward | CCGCTGCCGAAAATCTTTATCAACCAGCGTCTGGGC |
| Insert Reverse | GGTGGTGGTGGTGAAGCTTCGGTTGTTTGTTAATTTCATCGG |
| Backbone Forward | CCGATGAAATTAACAAACAACCGAAGCTTCACCACCACCACC |
| Backbone Reverse | GCCCAGACGCTGGTTGATAAAGATTTTCGGCAGCGG |

### Table of PCR Reaction Conditions and Thermocycling Parameters

**Table S3.** PCR reaction conditions and thermocycling parameters used to generate the SaNreA chimera plasmid.

**Insert Reaction Conditions**

| Component | Volume |
|---|---|
| Phusion Hot Start Flex 2X Master Mix | 25 µL |
| 10 ng/µL DNA Template | 1 µL |
| 10 µM Insert Forward Primer | 1 µL |
| 10 µM Insert Reverse Primer | 1 µL |
| DMSO | 2 µL |
| Autoclaved Water | 20 µL |
| **Total** | 50 µL |

**Insert Thermocycler Settings**

<table>
<tr><th>Step</th><th>Temperature</th><th>Time</th><th>Number of Cycles</th></tr>
<tr><td rowspan="2">Template Denaturation</td><td>98 °C</td><td>3 min</td><td>1</td></tr>
<tr><td>98 °C</td><td>30 sec</td><td rowspan="3">30</td></tr>
<tr><td>Annealing</td><td>55 °C</td><td>30 sec</td></tr>
<tr><td>Extension</td><td>72 °C</td><td>30 sec</td></tr>
<tr><td>Final Extension</td><td>72 °C</td><td>5 min</td><td>1</td></tr>
<tr><td>Storage</td><td>10 °C</td><td>∞</td><td>1</td></tr>
</table>

**Backbone Reaction Conditions**

| Component | Volume |
|---|---|
| Phusion Hot Start Flex 2X Master Mix | 25 µL |
| 10 ng/µL DNA Template | 1 µL |
| 10 µM Backbone Forward Primer | 1 µL |
| 10 µM Backbone Reverse Primer | 1 µL |
| DMSO | 2 µL |
| Autoclaved Water | 20 µL |
| **Total** | 50 µL |

**Backbone Thermocycler Settings**

<table>
<tr><th>Step</th><th>Temperature</th><th>Time</th><th>Number of Cycles</th></tr>
<tr><td rowspan="2">Template Denaturation</td><td>98 °C</td><td>3 min</td><td>1</td></tr>
<tr><td>98 °C</td><td>30 sec</td><td rowspan="3">30</td></tr>
<tr><td>Annealing</td><td>55 °C</td><td>30 sec</td></tr>
<tr><td>Extension</td><td>72 °C</td><td>4 min</td></tr>
<tr><td>Final Extension</td><td>72 °C</td><td>10 min</td><td>1</td></tr>
<tr><td>Storage</td><td>10 °C</td><td>∞</td><td>1</td></tr>
</table>

### Size Exclusion Chromatograms and SDS-PAGE Gels

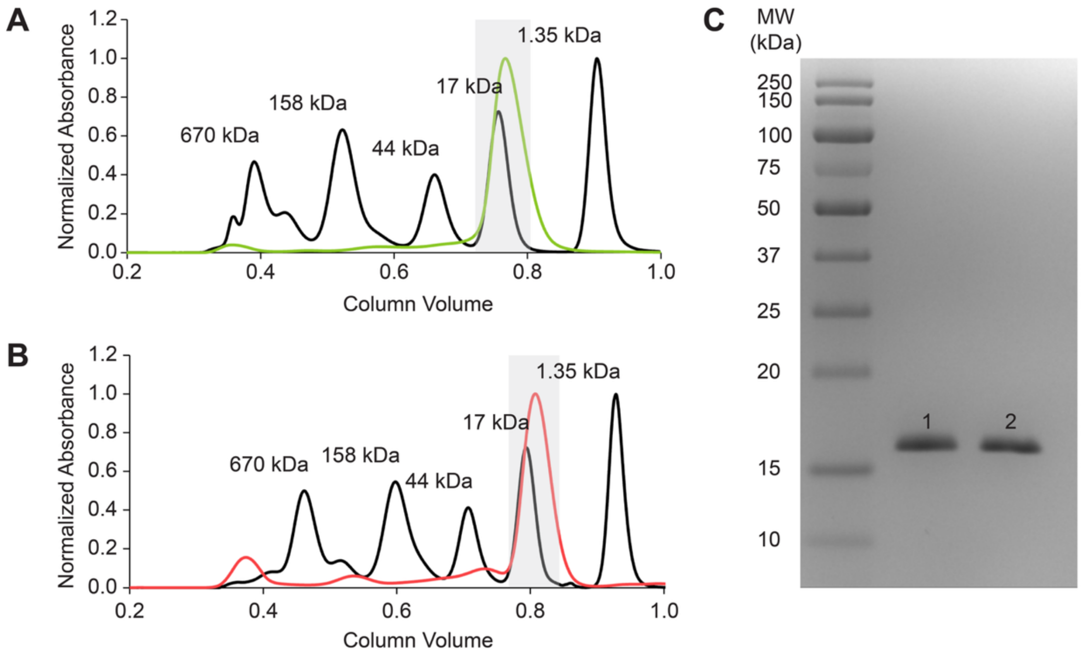

**Figure S15.** Normalized size-exclusion chromatograms monitored by absorbance at 280 nm for the (A) first protein preparation (green) and (B) second protein preparation (red) for SsimNreA with gel filtration standard (black) in 50 mM HEPES buffer (pH 7.8 at 10 ºC) with 200 mM NaCl. The gel filtration standard has thyroglobulin, γ-globulin, ovalbumin, myoglobin, and vitamin B12, shown from left to right. The gray-shaded region indicates the fractions that were collected. (C) Sodium dodecyl sulfate-polyacrylamide gel electrophoresis (SDS-PAGE) analysis of the isolated proteins. The protein ladder and purified SsimNreA are shown from left to right. The theoretical molecular weight of SsimNreA is approximately 18.4 kDa. Abbreviation: MW, molecular weight; kDa, kilodalton.

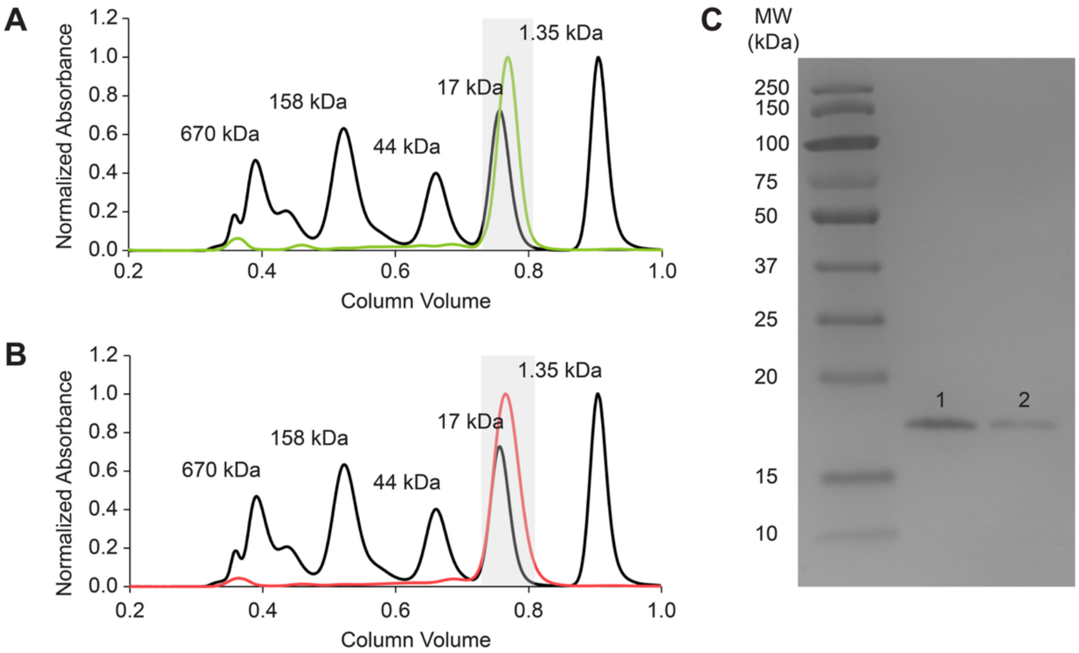

**Figure S16.** Normalized size-exclusion chromatograms monitored by absorbance at 280 nm for the (A) first protein preparation (green) and (B) second protein preparation (red) for ShNreA with gel filtration standard (black) in 50 mM HEPES buffer (pH 7.8 at 10 ºC) with 200 mM NaCl. The gel filtration standard has thyroglobulin, γ-globulin, ovalbumin, myoglobin, and vitamin B12, shown from left to right. The gray-shaded region indicates the fractions that were collected. (C) SDS-PAGE analysis of the isolated proteins. The protein ladder and purified ShNreA are shown from left to right. The theoretical molecular weight of ShNreA is approximately 18.4 kDa.

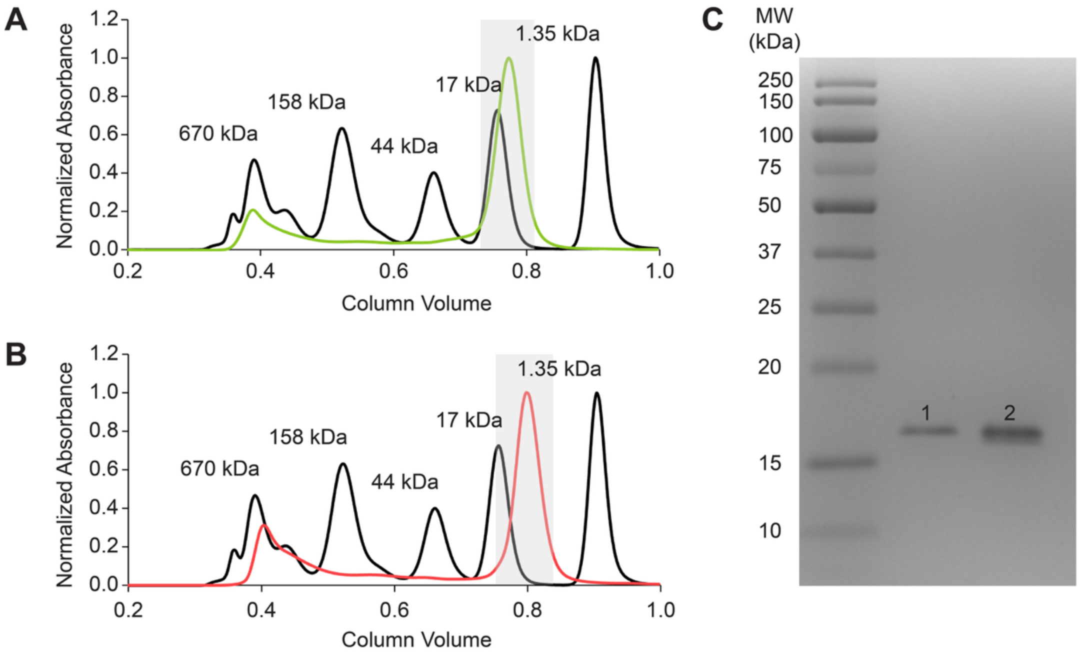

**Figure S17.** Normalized size-exclusion chromatograms monitored by absorbance at 280 nm for the (A) first protein preparation (green) and (B) second protein preparation (red) for SarNreA with gel filtration standard (black) in 50 mM HEPES buffer (pH 7.8 at 10 ºC) with 200 mM NaCl. The gel filtration standard has thyroglobulin, γ-globulin, ovalbumin, myoglobin, and vitamin B12, shown from left to right. The gray-shaded region indicates the fractions that were collected. (C) SDS-PAGE analysis of the isolated proteins. The protein ladder and purified SarNreA are shown from left to right. The theoretical molecular weight of SarNreA is approximately 18.7 kDa.

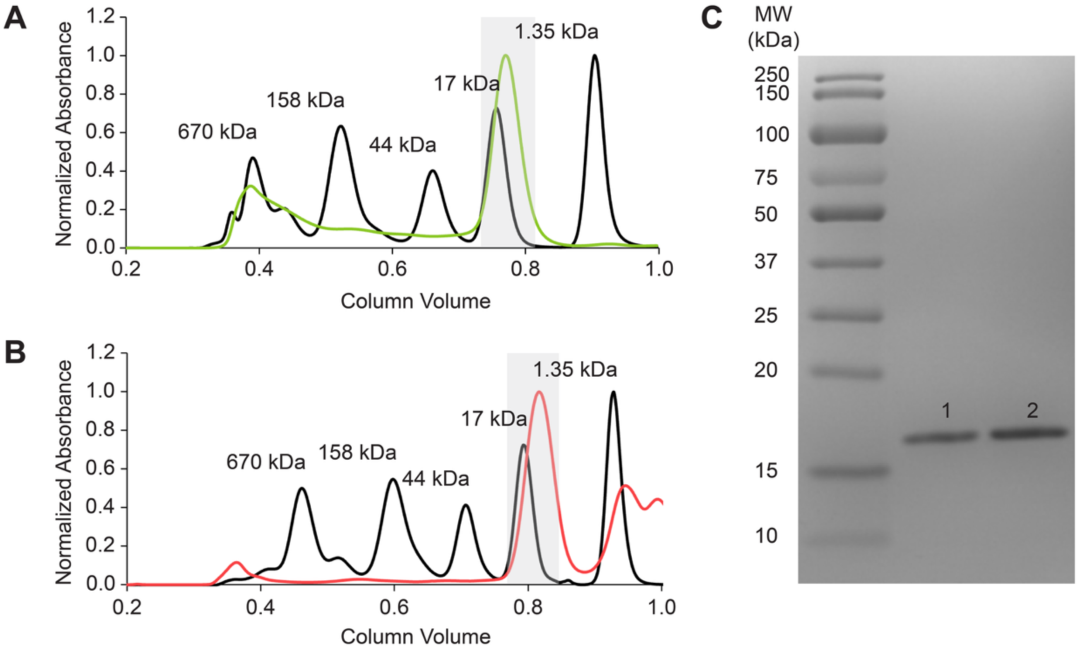

**Figure S18.** Normalized size-exclusion chromatograms monitored by absorbance at 280 nm for the (A) first protein preparation (green) and (B) second protein preparation (red) for SsNreA with gel filtration standard (black) in 50 mM HEPES buffer (pH 7.8 at 10 ºC) with 200 mM NaCl. The gel filtration standard has thyroglobulin, γ-globulin, ovalbumin, myoglobin, and vitamin B12, shown from left to right. The gray-shaded region indicates the fractions that were collected. (C) SDS-PAGE analysis of the isolated proteins. The protein ladder and purified SsNreA are shown from left to right. The theoretical molecular weight of SsNreA is approximately 18.7 kDa.

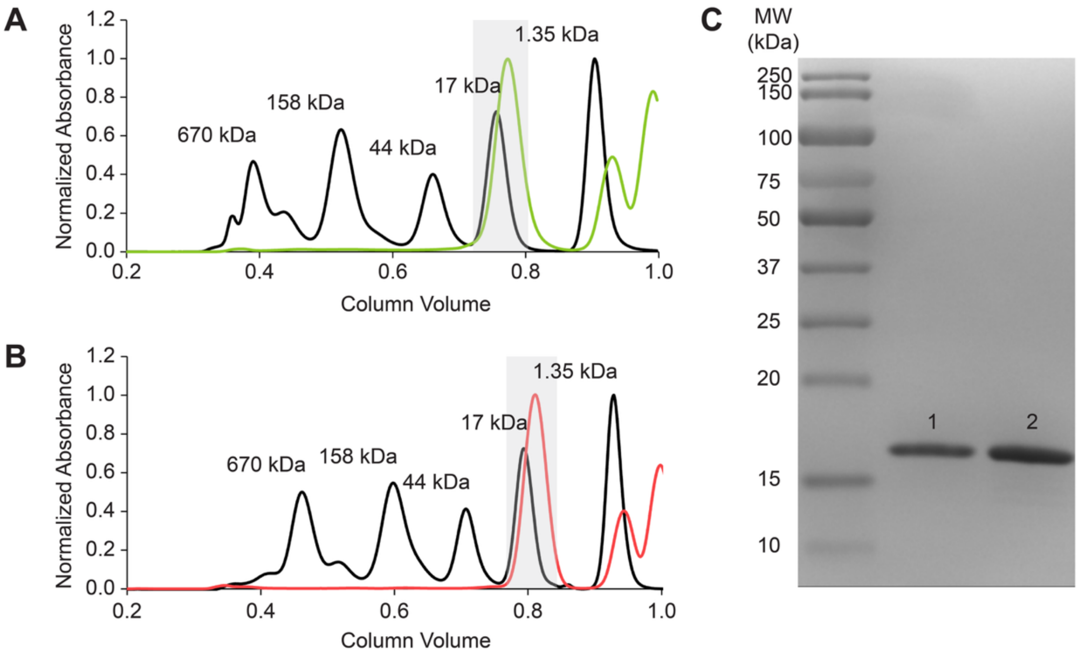

**Figure S19.** Normalized size-exclusion chromatograms monitored by absorbance at 280 nm for the (A) first protein preparation (green) and (B) second protein preparation (red) for SaNreA with gel filtration standard (black) in 50 mM HEPES buffer (pH 7.8 at 10 ºC) with 200 mM NaCl. The gel filtration standard has thyroglobulin, γ-globulin, ovalbumin, myoglobin, and vitamin B12, shown from left to right. The gray-shaded region indicates the fractions that were collected. (C) SDS-PAGE analysis of the isolated proteins. The protein ladder and purified SaNreA are shown from left to right. The theoretical molecular weight of SaNreA is approximately 18.2 kDa.

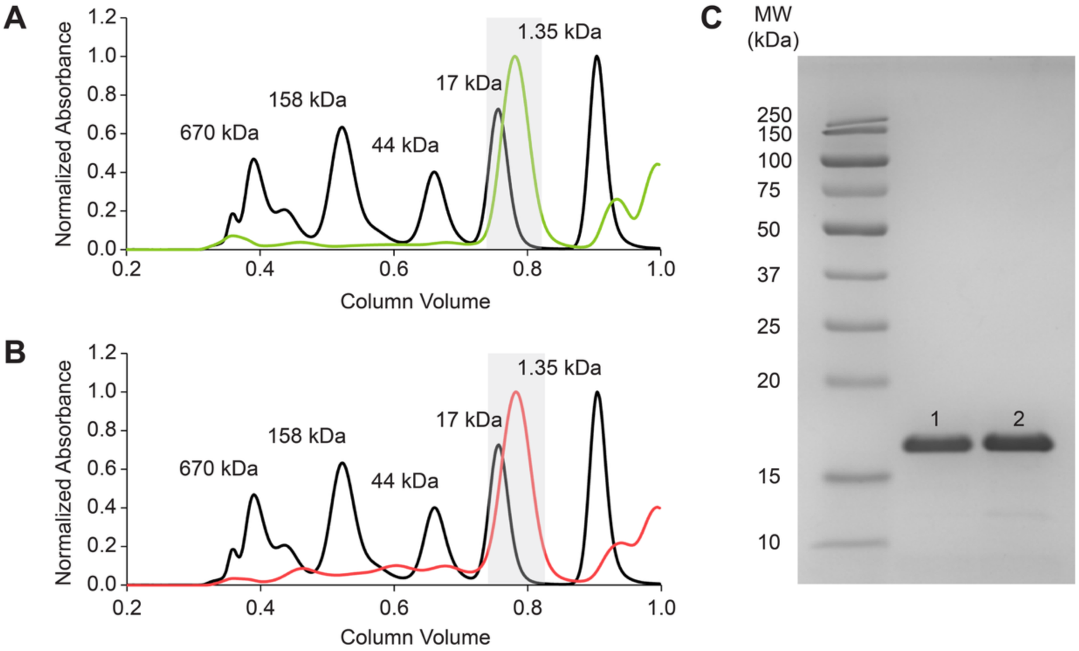

**Figure S20.** Normalized size-exclusion chromatograms monitored by absorbance at 280 nm for the (A) first protein preparation (green) and (B) second protein preparation (red) for the SaNreA chimera with gel filtration standard (black) in 50 mM HEPES buffer (pH 7.8 at 10 ºC) with 200 mM NaCl. The gel filtration standard has thyroglobulin, γ-globulin, ovalbumin, myoglobin, and vitamin B12, shown from left to right. The gray-shaded region indicates the fractions that were collected. (C) SDS-PAGE analysis of the isolated proteins. The protein ladder and purified SaNreA chimera are shown from left to right. The theoretical molecular weight of SaNreA chimera is approximately 18.1 kDa.

### Table of Theoretical Molar Extinction Coefficients

**Table S4.** Theoretical molar extinction coefficients for the proteins tested in this study.

| Protein | Molar Extinction Coefficient at 280 nm ($M^{-1}$ $cm^{-1}$) |
|---|---|
| ScNreA | 21430 |
| SsimNreA | 18450 |
| ShNreA | 22920 |
| SarNreA | 22920 |
| SsNreA | 19940 |
| SaNreA | 19940 |
| SaNreA chimera | 19940 |

### Thermodynamic Characterization: Thermograms and Isotherms

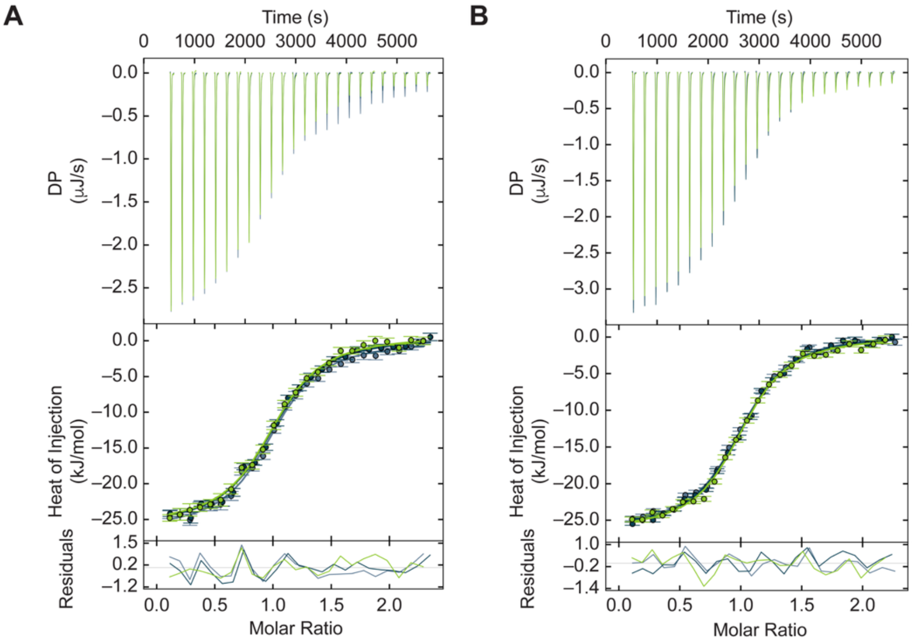

**Figure S21.** Analysis of nitrate binding to SsimNreA by ITC at 20 °C. The SVD-reconstructed thermogram, binding isotherm with error estimates for each injection, and residuals are shown for three technical replicates from the (A) first and (B) second protein preparations.

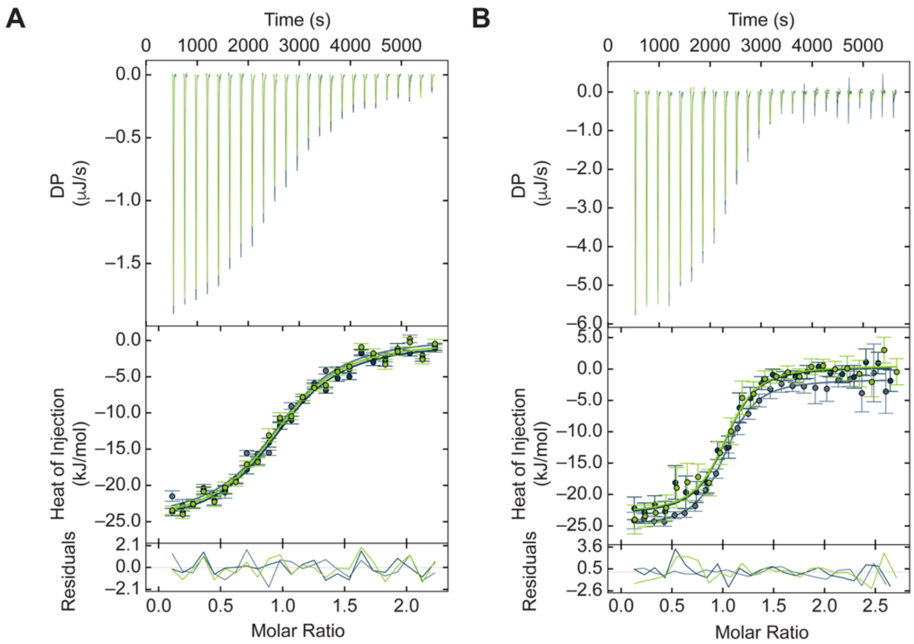

**Figure S22.** Analysis of nitrate binding to ShNreA by ITC at 20 °C. The SVD-reconstructed thermogram, binding isotherm with error estimates for each injection, and residuals are shown for three technical replicates from the (A) first and (B) second protein preparations.

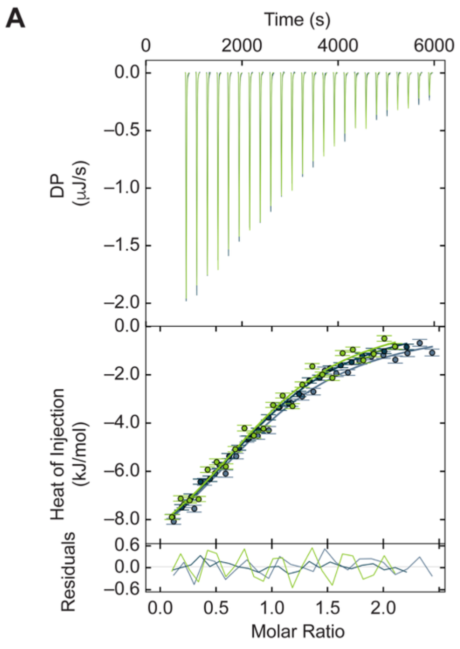

**Figure S23.** Analysis of nitrate binding to SarNreA by ITC at 20 °C. The SVD-reconstructed thermogram, binding isotherm with error estimates for each injection, and residuals are shown for three technical replicates from the (A) first and (B) second protein preparations.

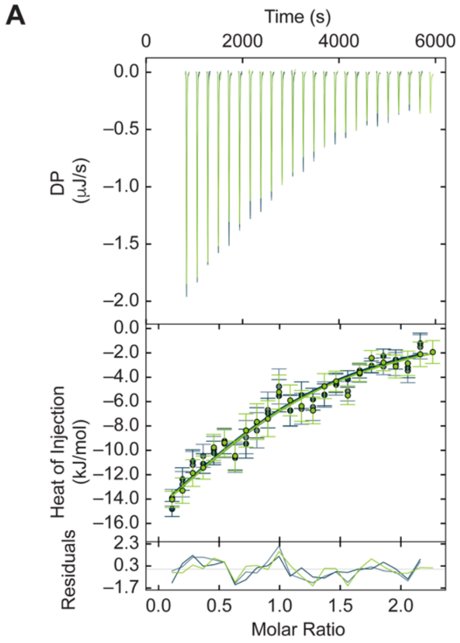

**Figure S24.** Analysis of nitrate binding to SsNreA by ITC at 20 °C. The SVD-reconstructed thermogram, binding isotherm with error estimates for each injection, and residuals are shown for three technical replicates from the (A) first and (B) second protein preparations.

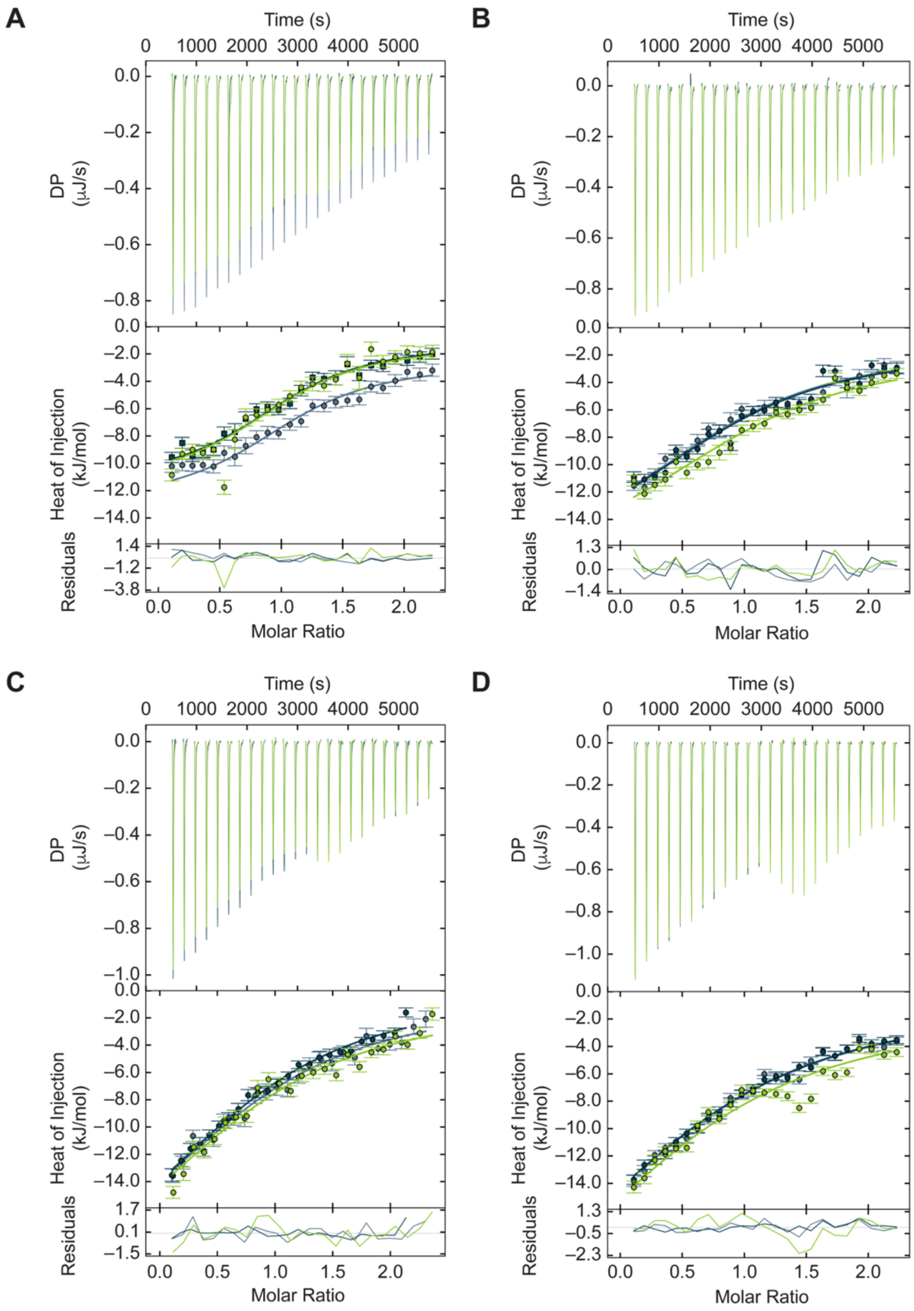

**Figure S25.** Analysis of nitrate binding to SaNreA by ITC at (A) 10 °C, (B) 15 °C, (C) 20 °C, and (D) 25 °C. The SVD-reconstructed thermogram, binding isotherm with error estimates for each injection, and residuals are shown for three technical replicates from the first protein preparation. Data for the second protein preparation is shown in Figure S26.

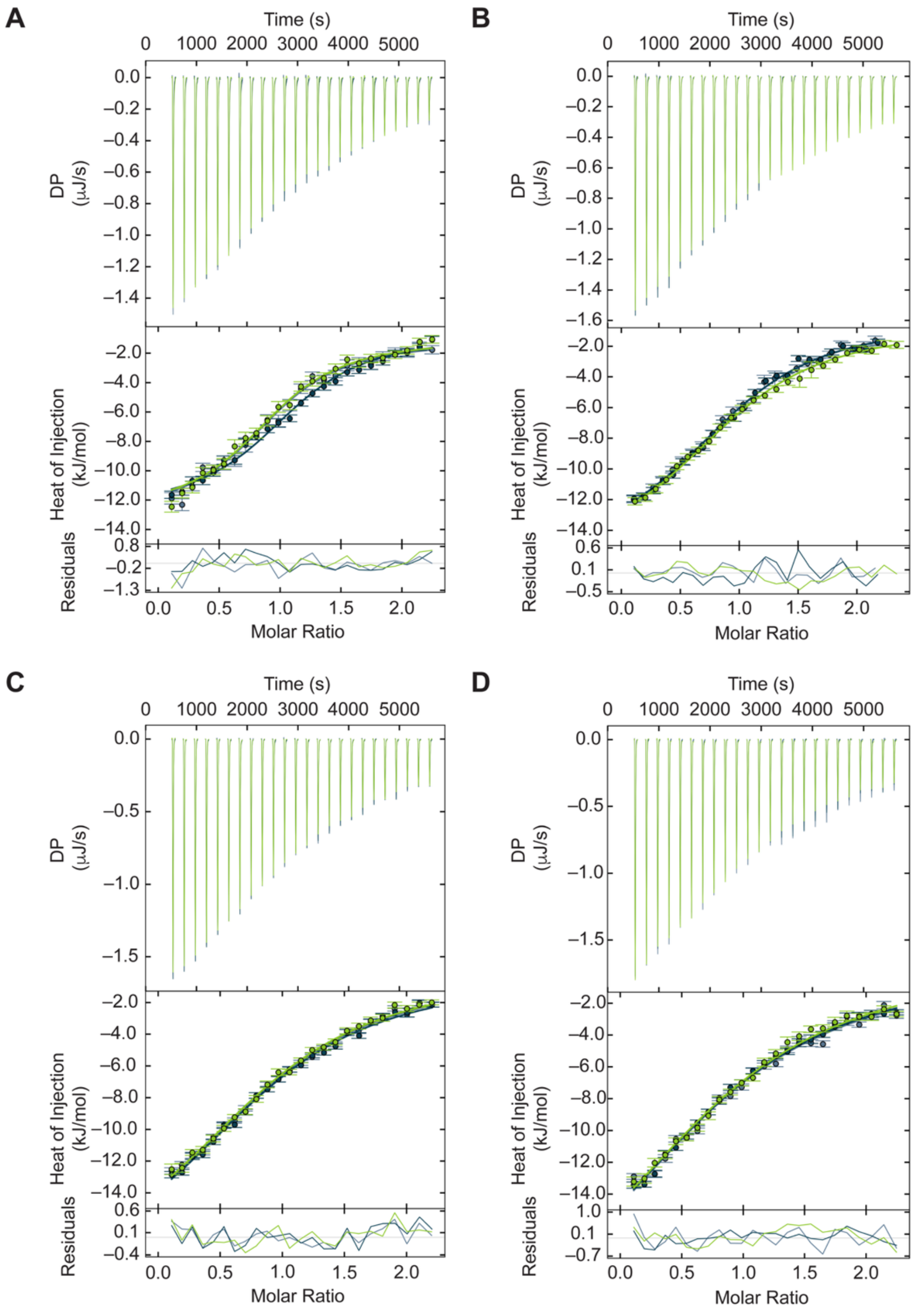

**Figure S26.** Analysis of nitrate binding to SaNreA by ITC at (A) 10 °C, (B) 15 °C, (C) 20 °C, and (D) 25 °C. The SVD-reconstructed thermogram, binding isotherm with error estimates for each injection, and residuals are shown for three technical replicates from the second protein preparation. Data for the first protein preparation is shown in Figure S25.

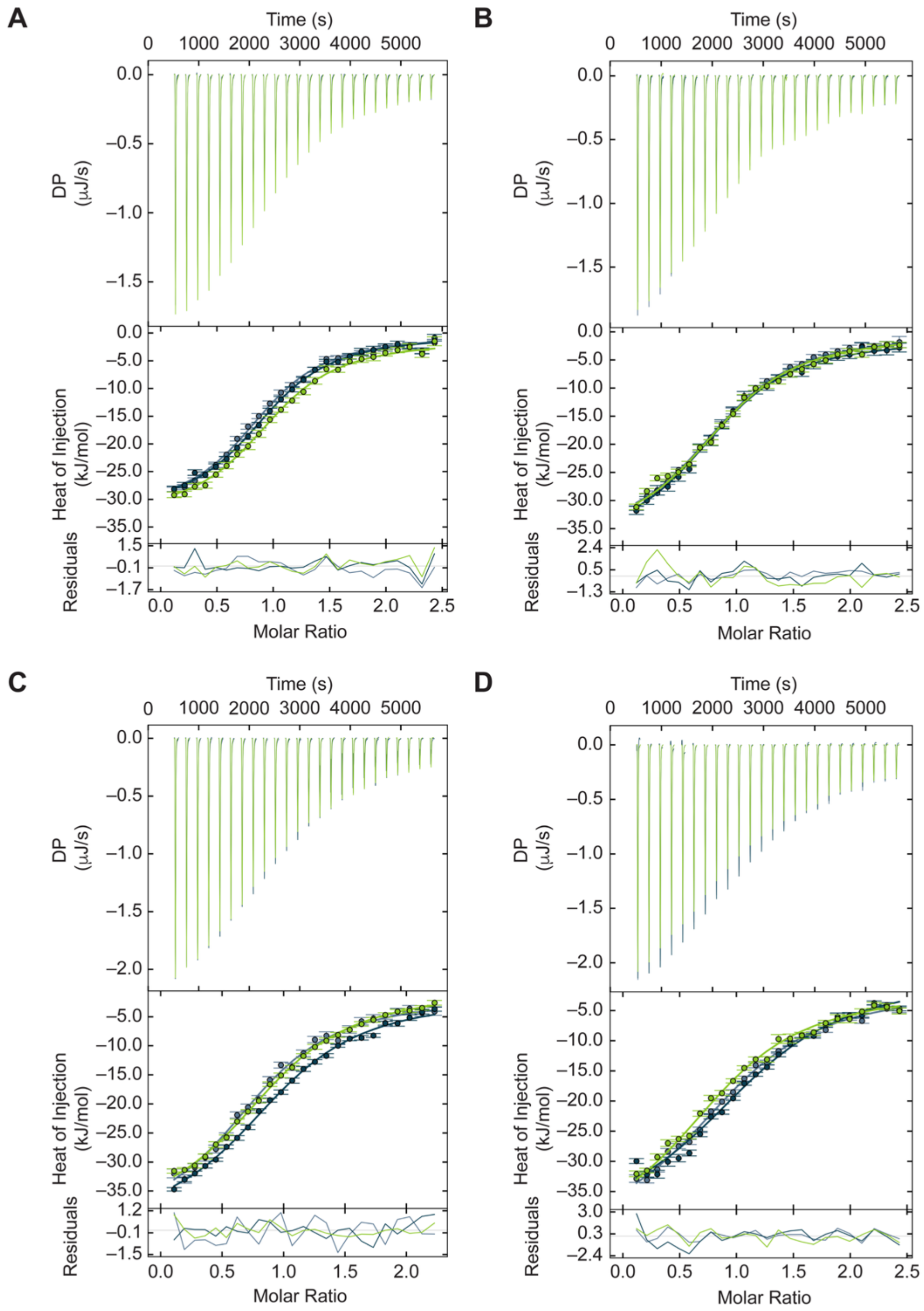

**Figure S27.** Analysis of nitrate binding to the SaNreA chimera by ITC at (A) 10 °C, (B) 15 °C, (C) 20 °C, and (D) 25 °C. The SVD-reconstructed thermogram, binding isotherm with error estimates for each injection, and residuals are shown for three technical replicates from the first protein preparation. Data for the second protein preparation is shown in Figure S28.

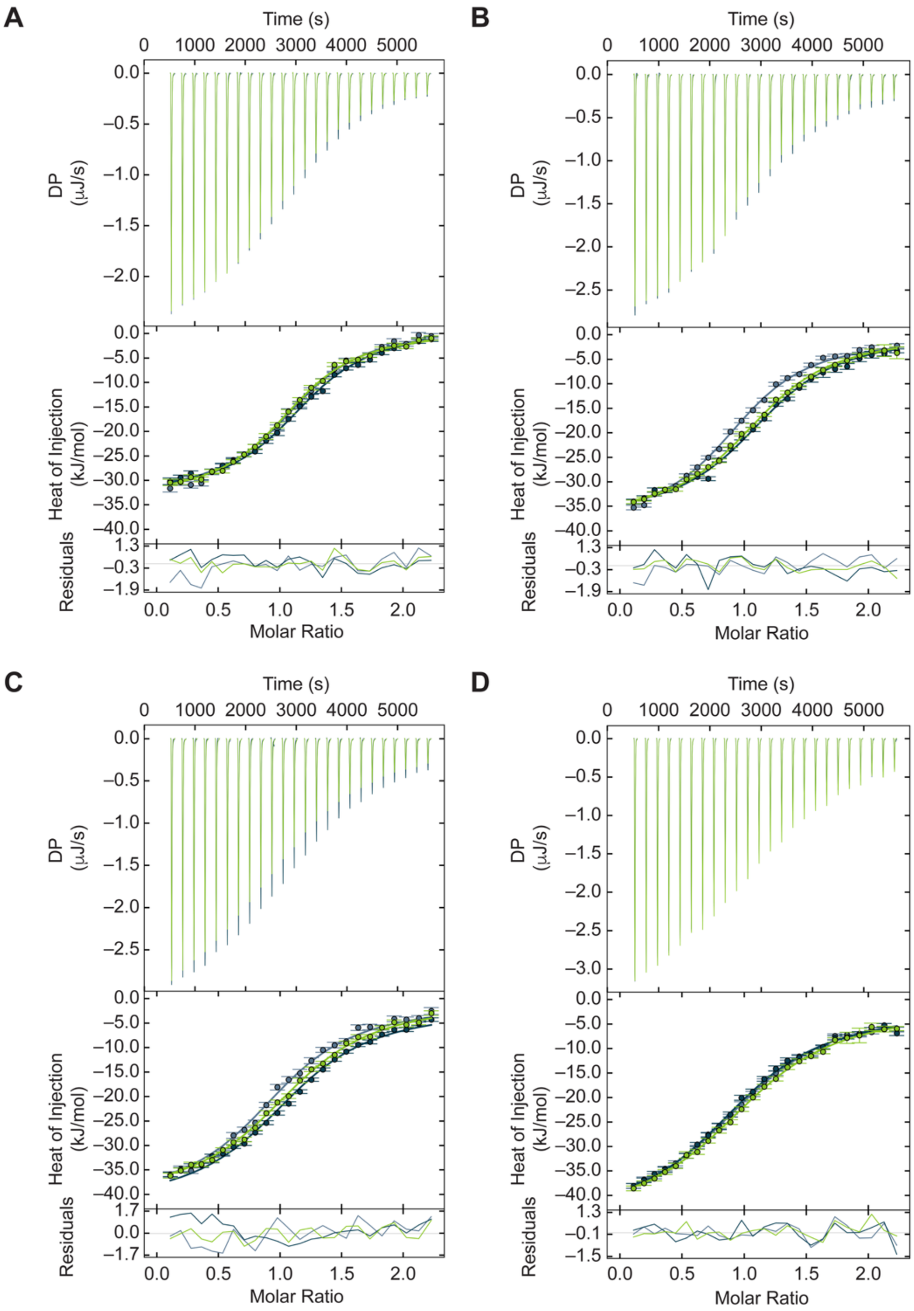

**Figure S28.** Analysis of nitrate binding to the SaNreA chimera by ITC at (A) 10 °C, (B) 15 °C, (C) 20 °C, and (D) 25 °C. The SVD-reconstructed thermogram, binding isotherm with error estimates for each injection, and residuals are shown for three technical replicates from the second protein preparation. Data for the first protein preparation is shown in Figure S27.

### Thermodynamic Characterization: Data Tables

**Table S5.** The thermodynamic parameters for SsimNreA binding to nitrate. The confidence intervals at 68.3% confidence are in brackets. The protein concentrations were 100 μM and 120 μM, and the stock nitrate concentrations were 1000 μM and 1200 μM for the first and second protein preparations, respectively.

| Protein Preparation | Temperature (°C) | $K_d$ (μM) | ΔH (kJ/mol) | TΔS (kJ/mol) | ΔG (kJ/mol) | Concentration Correction Factor |
|---|---|---|---|---|---|---|
| 1 | 20 | 4.7 [4.5, 5.0] | –26.3 [–26.4, –26.1] | 3.6 | –29.9 | 0.97 [0.96, 0.98]<br>0.95 [0.94, 0.96]<br>0.97 [0.96, 0.98] |
| 2 | 20 | 4.9 [4.6, 5.1] | –26.4 [–26.5, –26.3] | 3.5 | –29.8 | 0.98 [0.97, 0.99]<br>0.99 [0.98, 1.01]<br>1.02 [1.01, 1.03] |

**Table S6.** The thermodynamic parameters for ShNreA binding to nitrate. The confidence intervals at 68.3% confidence are in brackets. The protein concentrations were 70 μM and 200 μM, and the stock nitrate concentrations were 700 μM and 2000 μM for the first and second protein preparations, respectively.

| Protein Preparation | Temperature (°C) | $K_d$ (μM) | ΔH (kJ/mol) | TΔS (kJ/mol) | ΔG (kJ/mol) | Concentration Correction Factor |
|---|---|---|---|---|---|---|
| 1 | 20 | 5.1 [4.8, 5.3] | –25.2 [–25.4, –25.0] | 4.5 | –29.7 | 1.00 [0.98, 1.02]<br>1.00 [0.98, 1.02]<br>0.99 [0.97, 1.01] |
| 2 | 20 | 4.2 [3.9, 4.4] | –23.5 [–23.7, –23.4] | 6.7 | –30.2 | 0.94 [0.93, 0.95]<br>0.93 [0.90, 0.95]<br>0.90 [0.88, 0.93] |

**Table S7.** The thermodynamic parameters for SarNreA binding to nitrate. The confidence intervals at 68.3% confidence are in brackets. The protein concentrations were 240 μM and 160 μM, and the stock nitrate concentrations were 2400 μM and 1600 μM for the first and second protein preparations, respectively.

| Protein Preparation | Temperature (°C) | $K_d$ (μM) | ΔH (kJ/mol) | TΔS (kJ/mol) | ΔG (kJ/mol) | Concentration Correction Factor |
|---|---|---|---|---|---|---|
| 1 | 20 | 86.0 [83.9, 88.2] | –11.4 [–11.5, –11.3] | 11.4 | –22.8 | 0.95 [0.93, 0.98]<br>1.05 [1.03, 1.07]<br>1.10 [1.08, 1.13] |
| 2 | 20 | 103.1 [98.1, 108.3] | –11.9 [–12.0, –11.7] | 10.5 | –22.4 | 0.96 [0.88, 1.00]<br>0.97 [0.93, 1.00]<br>0.97 [0.93, 0.99] |

**Table S8.** The thermodynamic parameters for SsNreA binding to nitrate. The confidence intervals at 68.3% confidence are in brackets. The protein concentrations were 128 μM and 125 μM, and the stock nitrate concentrations were 1300 μM and 1250 μM for the first and second protein preparations, respectively.

| Protein Preparation | Temperature (°C) | $K_d$ (μM) | ΔH (kJ/mol) | TΔS (kJ/mol) | ΔG (kJ/mol) | Concentration Correction Factor |
|---|---|---|---|---|---|---|
| 1 | 20 | 97.2 [92.5, 102.1] | –27.1 [–27.4, –26.7] | –4.6 | –22.5 | 1.03 [1.00, 1.05]<br>1.01 [0.98, 1.03]<br>0.99 [0.96, 1.02] |
| 2 | 20 | 105.8 [103.3, 108.4] | –24.0 [–24.2, –23.9] | –1.7 | –22.3 | 1.08 [1.06, 1.10]<br>1.08 [1.06, 1.11]<br>1.08 [1.06, 1.10] |

**Table S9.** The thermodynamic parameters for SaNreA binding to nitrate. The confidence intervals at 68.3% confidence are in brackets. The protein concentration was 70 μM, and the stock nitrate concentration was 700 μM. Data for the second protein preparation is shown in Table S10.

| Temperature (°C) | $K_d$ (μM) | ΔH (kJ/mol) | TΔS (kJ/mol) | ΔG (kJ/mol) | Concentration Correction Factor |
|---|---|---|---|---|---|
| 10 | 12.7 [11.7, 13.9] | –10.2 [–10.4, –10.1] | 16.3 | –26.5 | 1.11 [1.07, 1.15]<br>1.06 [1.02, 1.14]<br>1.05 [1.01, 1.09] |
| 15 | 35.5 [33.7, 37.3] | –15.9 [–16.1, –15.7] | 8.7 | –24.6 | 1.00 [0.98, 1.03]<br>1.01 [0.99, 1.04]<br>1.13 [1.10, 1.15] |
| 20 | 51.3 [49.1, 51.6] | –23.1 [–23.4, –23.0] | 1.0 | –24.1 | 0.97 [0.94, 0.99]<br>1.04 [1.02, 1.07]<br>0.94 [0.92, 0.97] |
| 25 | 64.5 [62.9, 66.1] | –25.8 [–26.1, –25.6] | –1.9 | –23.9 | 1.00 [0.97, 1.02]<br>1.00 [0.97, 1.02]<br>0.98 [0.96, 1.00] |

**Table S10.** The thermodynamic parameters for SaNreA binding to nitrate. The confidence intervals at 68.3% confidence are in brackets. The protein concentration was 119 μM, and the stock nitrate concentration was 1200 μM. Data for the first protein preparation is shown in Table S9.

| Temperature (°C) | $K_d$ (μM) | ΔH (kJ/mol) | TΔS (kJ/mol) | ΔG (kJ/mol) | Concentration Correction Factor |
|---|---|---|---|---|---|
| 10 | 14.4 [13.7, 17.2] | –11.8 [–11.9, –11.7] | 14.6 | –26.4 | 0.95 [0.94, 0.98]<br>1.10 [1.08, 1.12]<br>0.95 [0.93, 0.98] |
| 15 | 31.4 [30.7, 32.2] | –15.2 [–15.3, –15.1] | 9.7 | –24.8 | 1.04 [1.03, 1.05]<br>1.03 [1.02, 1.04]<br>0.96 [0.95, 0.97] |
| 20 | 54.8 [53.6, 56.1] | –19.4 [–19.5, –19.3] | 4.5 | –23.9 | 1.02 [1.01, 1.03]<br>1.02 [1.01, 1.03]<br>1.02 [1.00, 1.03] |
| 25 | 81.2 [79.1, 82.9] | –24.7 [–24.8, –24.6] | –1.4 | –23.4 | 1.01 [0.99, 1.02]<br>1.01 [1.00, 1.02]<br>1.03 [1.01, 1.05] |

**Table S11.** The thermodynamic parameters for the SaNreA chimera binding to nitrate. The confidence intervals at 68.3% confidence are in brackets. The protein concentration was 55 μM, and the stock nitrate concentration was 550 μM. Data for the second protein preparation is shown in Table S12.

| Temperature (°C) | $K_d$ (μM) | ΔH (kJ/mol) | TΔS (kJ/mol) | ΔG (kJ/mol) | Concentration Correction Factor |
|---|---|---|---|---|---|
| 10 | 6.3 [6.2, 6.5] | –31.5 [–31.7, –31.4] | –3.4 | –28.2 | 0.92 [0.91, 0.93]<br>0.97 [0.96, 0.98]<br>1.01 [1.00, 1.02] |
| 15 | 9.4 [9.2, 9.6] | –36.7 [–36.9, –36.5] | –9.0 | –27.7 | 0.92 [0.91, 0.93]<br>0.89 [0.87, 0.92]<br>0.92 [0.91, 0.94] |
| 20 | 11.1 [11.0, 11.5] | –39.4 [–39.5, –39.2] | –11.5 | –27.8 | 0.94 [0.93, 0.95]<br>1.05 [1.04, 1.06]<br>0.94 [0.93, 0.95] |
| 25 | 14.1 [13.7, 14.4] | –41.2 [–41.4, –41.0] | –13.5 | –27.7 | 1.03 [1.02, 1.04]<br>1.19 [1.18, 1.20]<br>0.93 [0.91, 0.96] |

**Table S12.** The thermodynamic parameters for the SaNreA chimera binding to nitrate. The confidence intervals at 68.3% confidence are in brackets. The protein concentration was 75 μM, and the stock nitrate concentration was 750 μM. Data for the first protein preparation is shown in Table S11.

| Temperature (°C) | $K_d$ (μM) | ΔH (kJ/mol) | TΔS (kJ/mol) | ΔG (kJ/mol) | Concentration Correction Factor |
|---|---|---|---|---|---|
| 10 | 6.0 [5.9, 6.2] | –33.8 [–33.9, –33.7] | –5.5 | –28.3 | 1.15 [1.14, 1.16]<br>1.20 [1.19, 1.21]<br>1.11 [1.10, 1.12] |
| 15 | 7.3 [7.2, 7.5] | –37.0 [–37.1, –36.8] | –8.6 | –28.3 | 0.99 [0.98, 1.00]<br>1.18 [1.17, 1.19]<br>1.14 [1.13, 1.15] |
| 20 | 9.6 [9.4, 9.9] | –39.5 [–39.7, –39.3] | –11.4 | –28.0 | 1.03 [1.01, 1.05]<br>1.12 [1.11, 1.13]<br>1.13 [1.12, 1.14] |
| 25 | 12.1 [11.8, 12.4] | –42.2 [–42.3, –42.1] | –14.1 | –28.1 | 1.06 [1.05, 1.07]<br>1.02 [1.01, 1.03]<br>1.09 [1.08, 1.10] |

**Table S13.** The change in heat capacity ($\Delta C_p$) for SaNreA binding to nitrate. The confidence intervals at 68.3% confidence are in brackets. The protein concentrations were 70 μM and 119 μM, and the stock nitrate concentrations were 700 μM and 1200 μM for the first and second protein preparations, respectively.

| Protein Preparation | $\Delta C_p$ (J/mol·K) |
|---|---|
| 1 | –828.9 [–871.1, –787.0] |
| 2 | –638.1 [–679.9, –595.8] |

**Table S14.** The $\Delta C_p$ for the SaNreA chimera binding to nitrate. The confidence intervals at 68.3% confidence are in brackets. The protein concentrations were 55 μM and 75 μM, and the stock nitrate concentrations were 550 μM and 750 μM for the first and second protein preparations, respectively.

| Protein Preparation | $\Delta C_p$ (J/mol·K) |
|---|---|
| 1 | –632.2 [–674.0, –589.9] |
| 2 | –571.1 [–613.4, –529.3] |

### Circular Dichroism Spectra

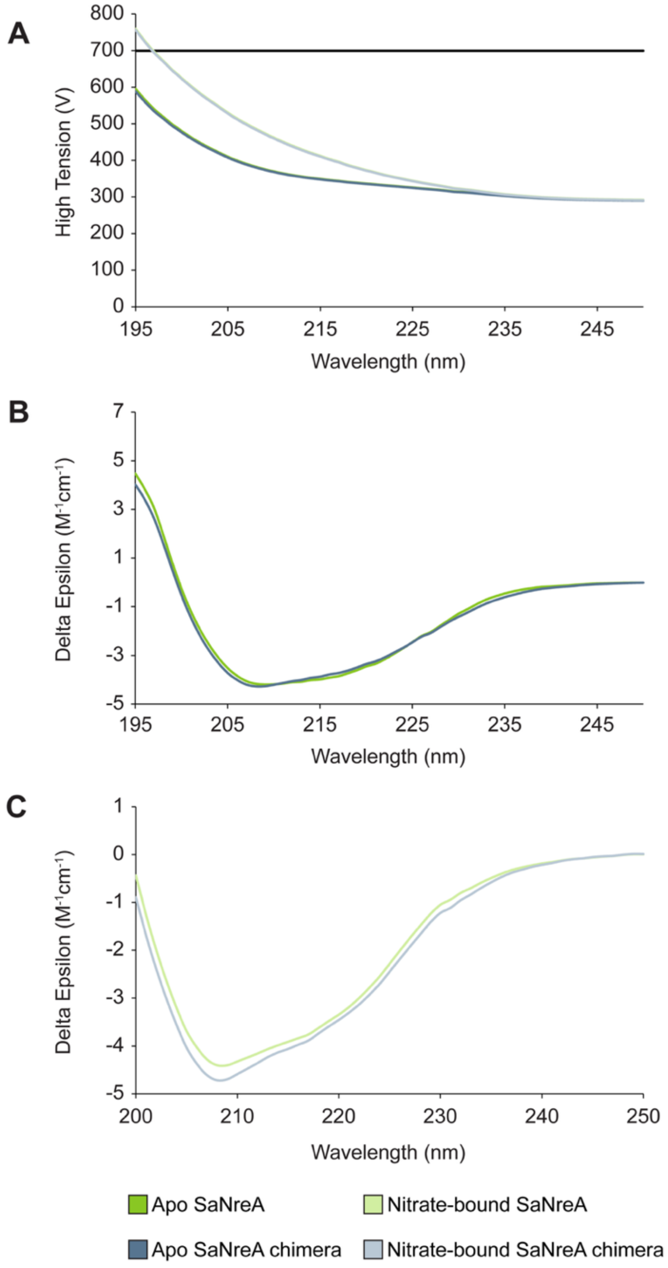

**Figure S29.** Circular dichroism (CD) characterization of apo and nitrate-bound SaNreA and the SaNreA chimera from the first protein preparation. (A) High-tension voltage as a function of wavelength, with the black line indicating 700 V. CD spectra for (B) apo and (C) nitrate-bound SaNreA and the SaNreA chimera. Spectra were collected using 5.5 μM SaNreA or SaNreA chimera in the absence or presence of 1 mM nitrate. Data for the second protein preparation is shown in Figure S30.

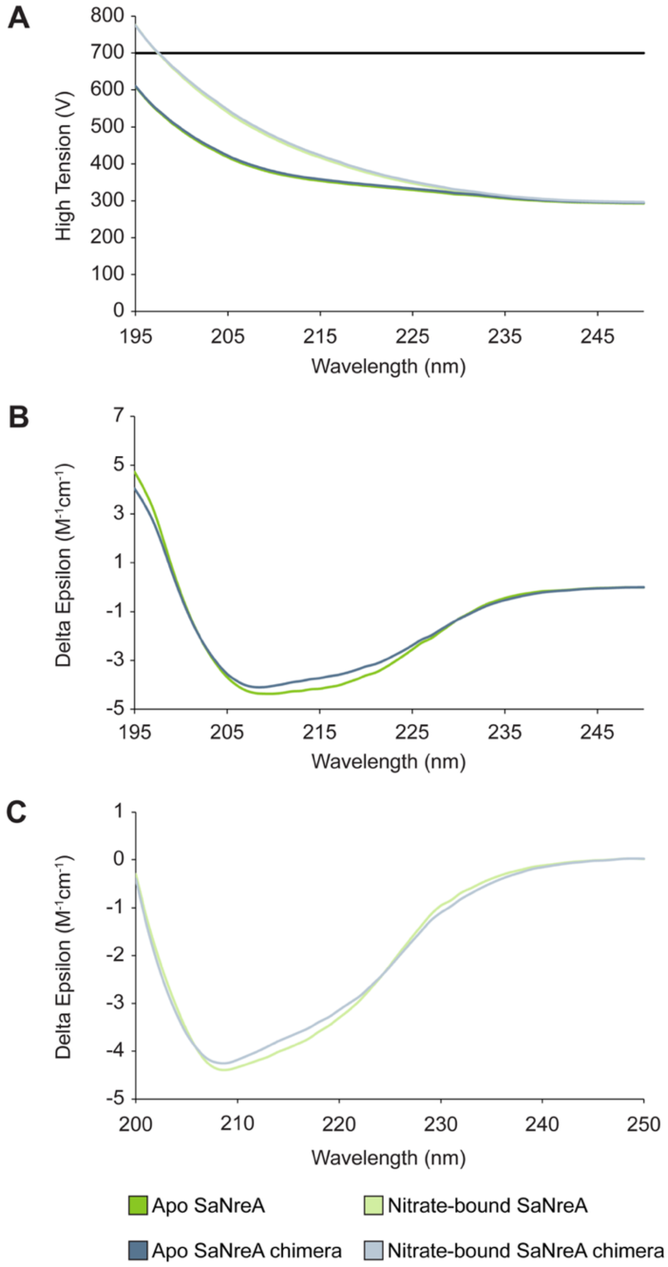

**Figure S30.** CD characterization of apo and nitrate-bound SaNreA and the SaNreA chimera from the second protein preparation. (A) High-tension voltage as a function of wavelength, with the black line indicating 700 V. CD spectra for (B) apo and (C) nitrate-bound SaNreA and the SaNreA chimera. Spectra were collected using 5.5 μM SaNreA or SaNreA chimera in the absence or presence of 1 mM nitrate. Data for the first protein preparation is shown in Figure S29.

### C-Terminal Interaction Networks

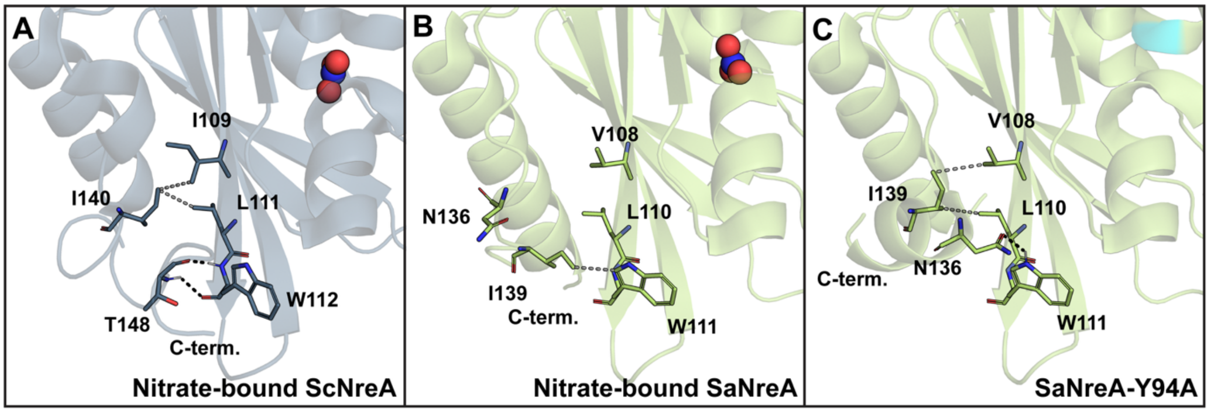

**Figure S31.** Polar (black dashed lines) and hydrophobic (gray dashed lines) interactions for (A) nitrate-bound ScNreA (PDB ID: 4IUK), (B) nitrate-bound SaNreA (PDB ID: 6IZJ), and (C) SaNreA-Y94A (PDB ID: 6K2H). The EDO molecules are omitted for clarity. The PyMOL "action: find: polar interactions" tool and measurement tool were used to find all polar and hydrophobic interactions within 4 Å.

### Back Views of AlphaFold3 Models with Confidence Levels

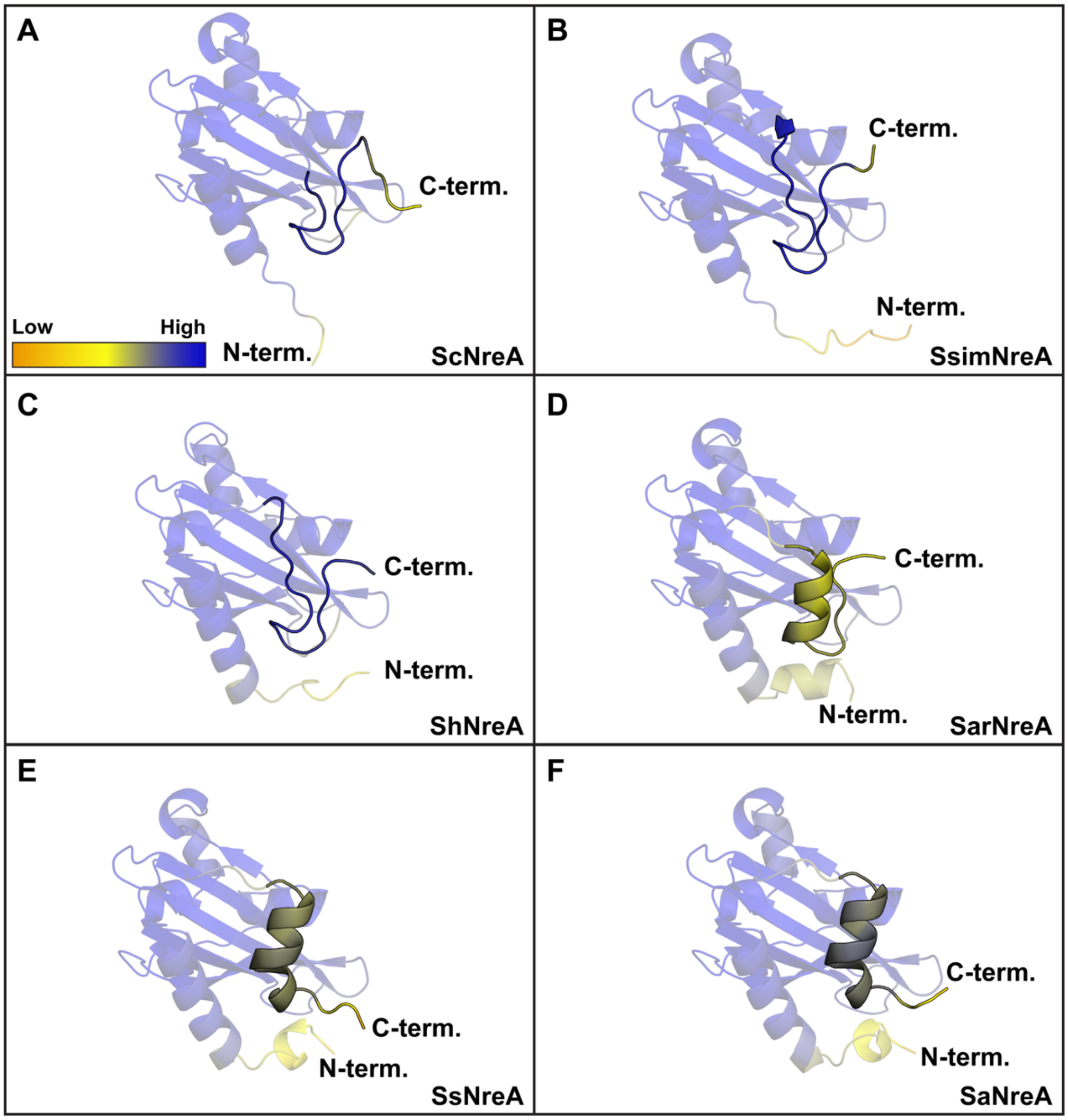

**Figure S32.** Back views of the AlphaFold3 models highlighting the predicted model confidence and C-terminal structural differences for the high-affinity NreAs, (A) ScNreA, (B) SsimNreA, and (C) ShNreA, and the low-affinity NreAs, (D) SarNreA, (E) SsNreA, and (F) SaNreA. Dark blue represents high confidence, and bright orange represents low confidence. Residues M1–I6 of ScNreA are not shown for clarity at the displayed figure scale.

### AlphaFold3 Models with Confidence Levels for the ScNreA and SaNreA Chimeras

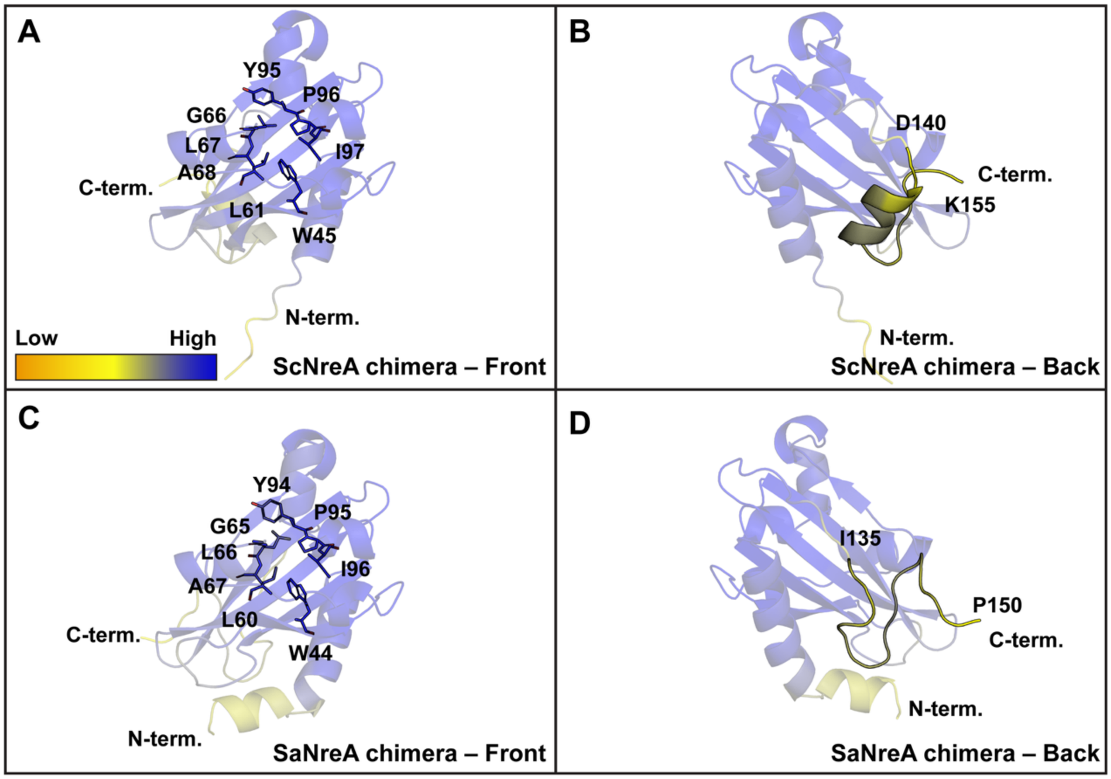

**Figure S33.** (A) Front and (B) back views of the AlphaFold3 model of the ScNreA chimera and (C) front and (D) back views of the AlphaFold3 model of the SaNreA chimera. The front views show the nitrate-binding pocket residues, and the back views highlight the predicted C-terminal conformations. Residues M1–I6 of the ScNreA chimera are not shown for clarity at the displayed figure scale.

### Summary Table

**Table S15.** C-terminal information for the NreA homologues and chimera tested in this study.

| Protein | C-Terminal Sequence | Overall Sequence Identity to ScNreA (%) | C-Terminal Sequence Identity to ScNreA (%) | AlphaFold3 Predicted C-Terminal Structure | Affinity Group |
|---|---|---|---|---|---|
| ScNreA | INQRLGSFTDEINKQP | – | – | Loop-like | High |
| SsimNreA | EIRINHRLGSFRDEIN | 77.5 | 84.6 | Loop-like | High |
| ShNreA | ETLDISGKLWTFSEEM | 51.0 | 33.3 | Loop-like | High |
| SarNreA | DYEDIQYKFGIFNDDK | 47.3 | 33.3 | α-helix | Low |
| SsNreA | DNDDIQRKFGIFNDDK | 46.0 | 33.3 | α-helix | Low |
| SaNreA | DNDDIQRKFGIFNDDK | 46.0 | 33.3 | α-helix | Low |
| SaNreA chimera | INQRLGSFTDEINKQP | 54.0 | 100.0 | Loop-like | Intermediate |

### Genome Neighborhood Diagrams for Representative NreA Homologues

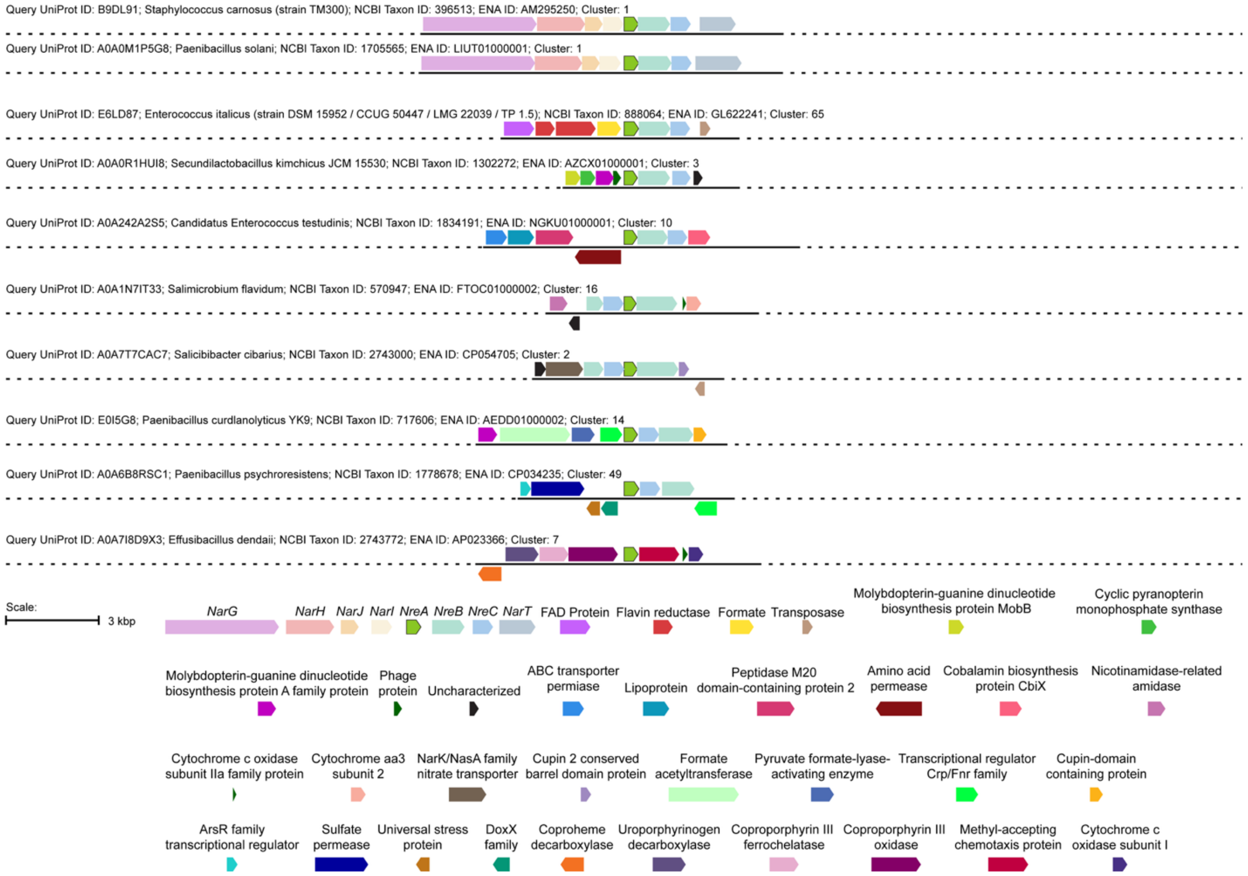

**Figure S34.** Genome neighborhood diagrams for ScNreA and representative NreA homologues identified through the phmmer search. For each diagram, the *nreA* gene (green) is highlighted along with neighboring genes encoding proteins with diverse predicted functions.
